# Pharmacological modulation of developmental and synaptic phenotypes in human SHANK3 deficient stem cell-derived neuronal models

**DOI:** 10.1101/2023.09.13.557523

**Authors:** Amandine Thibaudeau, Karen Schmitt, Louise François, Laure Chatrousse, David Hoffmann, Loïc Cousin, Amélie Weiss, Aurore Vuidel, Christina B Jacob, Peter Sommer, Alexandra Benchoua, Johannes H Wilbertz

## Abstract

**One sentence summary:** This study describes the use of SHANK3 deficient stem cell-derived neuronal models to screen and characterize small molecules that partially rescued developmental and synaptic defects related to Phelan-McDermid syndrome (PMDS).

Phelan-McDermid syndrome (PMDS) arises from mutations in the terminal region of chromosome 22q13, impacting the *SHANK3* gene. The resulting deficiency of the postsynaptic density scaffolding protein SHANK3 is associated with autism spectrum disorder (ASD). We examined 12 different PMDS patient and CRISPR-engineered stem cell-derived neuronal models and controls and found that reduced expression of SHANK3 leads to neuronal hyperdifferentiation, increased synapse formation, and decreased neuronal activity. We performed automated imaging-based screening of 7,120 target-annotated small molecules and identified three compounds that rescued SHANK3-dependent neuronal hyperdifferentiation. One compound, Benproperine, rescued the decreased colocalization of Actin Related Protein 2/3 Complex Subunit 2 (ARPC2) with ß-actin and rescued increased synapse formation in SHANK3 deficient neurons when administered early during differentiation. Neuronal activity was only mildly affected, highlighting Benproperine’s effects as a neurodevelopmental modulator. This study demonstrates that small molecular compounds that reverse developmental phenotypes can be identified in human neuronal PMDS models.

## Introduction

Phelan-McDermid syndrome (PMDS) is a rare genetic disorder characterized by developmental delay, hypotonia, intellectual disability (ID), speech impairments and autism spectrum disorder (ASD) (1–3). In most PMDS affected individuals the terminal region of chromosome 22q13 is impacted by point mutations or larger deletions. The gene SH3 and multiple ankyrin repeat domains 3 (*SHANK3*) is affected in most cases, often by larger deletions or frameshift truncations in exon 21 (4,5). *SHANK3* encodes a protein of the postsynaptic density (PSD) of excitatory synapses and likely fulfills a scaffolding function for the PSD complex. Consequently, synaptic roles of SHANK3 in PMDS have been studied extensively. PMDS neurons show for example reduced expression of glutamate receptors resulting in impaired excitatory synaptic transmission (6). Although SHANK3 is a primarily synaptic protein, SHANK3 deficiency has been shown to also have developmental consequences, for example by downregulating the proliferation of neural progenitor cells (NPCs) (7). Additionally, hyperdifferentiation related defects have been detected in SHANK3-deficient neurons. For example, the number of neurites per neuron was increased after 30 days of differentiation in patient iPSC-derived neurons while in Shank3B^−/−^ mice earlier maturation and hyperactivity of corticostriatal circuits has been observed (8,9).

Developmental mechanisms rely on neural activity for proper cortical development and equally, developmental mechanisms impact neural activity (10). Since NPCs reside in stem cell niches throughout life and contribute to adult neurogenesis, the possibility to modulate PMDS-related developmental and synaptic phenotypes in the postnatal brain might exist (11–14). For example, the activation of Shank3 expression positively impacted neural function and rescued certain autistic-like phenotypes in adult mice (15). Additionally, previous work has shown that SHANK3-related phenotypes can be rescued by pathway modulation of CLK2, ERK2 or direct upregulation of SHANK3 (16–20). First preclinical and clinical studies in SHANK3 human neuronal models or PMDS patient cohorts have begun to evaluate protein or small molecular compounds that normalize synaptic transmission or increase SHANK3 expression. PMDS-related synaptic defects were for example rescued with insulin-like growth factor 1 (IGF1) (6,21). Using a qPCR-based compound screening, Darville *et al.* identified Lithium as a compound positively impacting SHANK3 expression (17).

Despite these advances, studies combining both the pharmacological rescue of developmental and synaptic phenotypes in a SHANK3 deficiency context have not been performed. The dysregulation of both developmental and synaptic neurobiology needs to be better understood and eventually corrected to design a successful PMDS treatment strategy. Here we describe a hyperdifferentiation phenotype in PMDS-patient derived NPCs and reproduced these findings in CRISPR/Cas9-engineered SHANK3 deficient NPCs. CRISPR-engineered NPCs were then subjected to small molecular compound screening using a chemical library with known modes-of-action (MoA). We identified three SHANK3-defficiency specific small molecules able to rescue the hyperdifferentiation phenotype in both patient-derived and CRISPR-engineered NPCs. The compound Benproperine also rescued synapse formation but only mildly affected neuronal activity, highlighting its effect during neurodevelopment. Benproperine is an actin-related protein 2/3 complex subunit 2 (ARPC2) inhibitor. Treated cells presented a rescued ß-actin/ARPC2 colocalization phenotype. We anticipate that the use of chemical genomics approaches in human PMDS model systems can lead to the discovery and testing of cellular pathways that positively influence both cortical development and synaptic homeostasis which can increase the chances for the successful development of treatments against PMDS.

## Results

### SHANK3 haploinsufficiency results in hyperdifferentiation in a PMDS patient-derived cell line

Based on previous studies that identified signs of hyperdifferentiation in SHANK3 mutated PMDS models, we asked the question whether decreased proliferation and increased differentiation could be observed as early as in the NPC stage (7–9). We used three previously described human iPSC-based models to analyze the effects of SHANK3 *de novo* heterozygous truncating mutations on neuronal differentiation (ASD01, ASD03, ASD04). The donors presented with ASD and moderate to severe ID (4,17,22). All three mutations likely result in truncated SHANK3 by premature stop codons, either by direct mutation (E809X) or via frameshifting (G1271Afs*15 and L1142Vfs*153) (**Figure 1A**). Additionally, we used three healthy donor iPSC lines (PDF01, PC056, 4603). iPSCs were pre-differentiated for 20 days into NPCs and cryopreserved. After thawing, NPCs were differentiated into cortical neurons in 384-well plates and stained with Hoechst, and antibodies against the proliferation marker Ki67, and the neuronal marker HuC/D. Staining was performed at differentiation day 6 (D6), D9 and D14 (**Figure 1B**). We performed automated fluorescence confocal microscopy and image segmentation to quantify the ratios of proliferating and differentiating cells across all patient-derived cell lines. Compared to healthy donor cells, we observed decreased proliferation in ASD04 NPCs at D9 and D14. In line with decreased proliferation, also the ratio of HuC/D-positive cells was > 2-fold increased across all timepoints in ASD04 NPCs, indicating neuronal hyperdifferentiation (**Figure 1C**).

**Figure 1:**
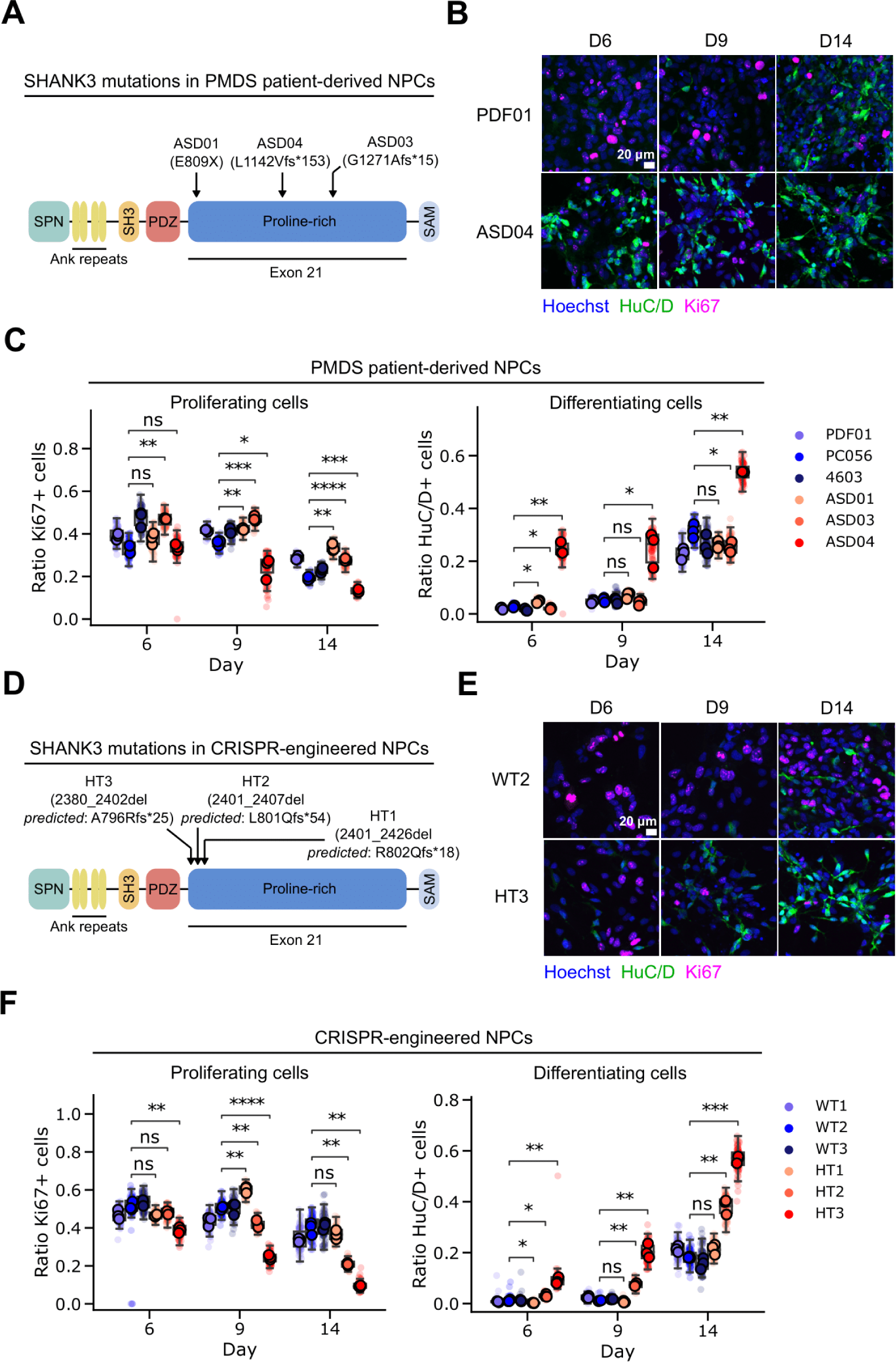
SHANK3-deficiency results in hyperdifferentiation in PMDS patient-derived and CRISPR-engineered cell lines. (A) Schematic depiction of SHANK3 domains and localization of mutations in the PMDS patient-derived iPSCs. fs* indicates a frame shift and gives the position of the stop codon in the new reading frame. (B) Fluorescently stained patient-derived NPCs at multiple differentiation timepoints. (C) Immunostaining-based quantification of the ratio of proliferating and differentiating healthy donor (blues) and patient-derived NPCs (reds). (D) Same as in (A), but mutations in the CRISPR-engineered cell lines are indicated. Frameshifts are predictions based on the observed deletion. (E) Same as (B), but in CRISPR-engineered isogenic NPC pairs. (F) Same as (C), but in CRISPR-engineered isogenic NPC pairs. Each datapoint represents technical replicate data from one 384-well plate well. Larger markers represent the medians of independent differentiation experiments. The boxplots show the median and the 2^nd^ and 3^rd^ quartiles of the data. Whiskers indicate 1.5X of the interquartile range. Welch’s unequal variances t-test was performed using the medians of independent differentiation experiments. ns = not significant, * = p < 0.05, ** = p < 0.01, *** = p < 0.001.

### SHANK3 haploinsufficiency results in hyperdifferentiation in CRISPR-engineered cell lines

Next, we wanted to confirm the hyperdifferentiation phenotype in unrelated SHANK3 CRISPR-engineered cells. The objective was to reproduce the loss of function mutations in the *SHANK3* gene in a heterozygous manner to model the mutations found in the patient cells. Since *SHANK3* mutations in PMDS patients are predominantly localized in exon 21, we used CRISPR/Cas9-engineered SHANK3 haploinsufficiency models generated with cut-sites in exon 21 in the human embryonic stem cell (hESC) line SA001. CRISPR-engineered clones were recovered, and the type of deletion analyzed using PCR and sequencing. All successful deletions were localized in the N-terminal part of *SHANK3’s* proline-rich domain (HT1, HT2, HT3) (**Figure 1D**). All deletions were predicted to lead to frameshifts and premature stop codons. gRNA-transfected, but unmutated clones were selected as isogenic control lines (WT1, WT2, WT3). Reduced levels of SHANK3 expression in the CRISPR-modified cells were confirmed by RT-qPCR and Western blotting (**Figure S1A-C**). Identical to the previous experiments in patient-derived NPCs, we stained the CRISPR-engineered NPCs at D6, D9, and D14 using antibodies against Ki67 and HuC/D (**Figure 1E**). Image quantification showed that 2 out of 3 CRISPR-engineered NPC lines showed a similar hyperdifferentiation phenotype as patient-derived cell line ASD04. CRISPR-engineered line HT3 showed decreased proliferation and increased neuronal differentiation across all tested days. HT2 presented a similar hyperdifferentiation phenotype on D9 and D14 (**Figure 1F**). In addition to Ki67 as proliferation marker, we also used an EdU (5-ethynyl-2’-deoxyuridine) incorporation assay. EdU becomes fluorescent when incorporated into replicating DNA, indicating cellular proliferation. 38% of WT2 and 8% of HT3 NPCs were EdU positive, corresponding to the values we obtained using the Ki67 staining (**Figure S1D**, **Figure 1F**). We also noted that the CRISPR-engineered isogenic control lines (WT1, WT2, WT3) and the healthy donor lines (PDF01, PC056, 4603) behaved homogenously and differentiated similarly, suggesting only small clonal or differentiation protocol induced variability. The observation that only two out of the three CRISPR-engineered NPC lines showed the hyperdifferentiation phenotype could be related to the slightly different SHANK3 truncation sites (**Figure 1D**). In summary, we used 12 mutant and control cell lines of different origins and observed that SHANK3 deficiency correlates in some cases with a hyperdifferentiation phenotype.

### SHANK3 haploinsufficiency results in altered synapse formation and decreased neuronal activity

To examine the effects of SHANK3-related hyperdifferentiation in neurons, we continued the differentiation of CRISPR-engineered lines WT2 and HT3 and patient-derived cell lines PC056 and ASD04 until 30 days of differentiation (D30). D30 neurons were stained using Hoechst and antibodies against SHANK3, Synapsin 1, and MAP2. To quantify the neuronal differentiation in all acquired images in an unbiased manner, we developed a computational image analysis workflow. All fluorescent image channels were segmented, and binary masks were used to quantify MAP2 network length or the position of synapses-related signals. To increase the confidence that detected fluorescent puncta indeed correspond to synapses we used several criteria: First, we searched for local maxima of SHANK3 and Synapsin1 spots and adpplied signal over noise, size and intensity thresholds. Secondly, we required both signals to be colocalized and to be localized to a MAP2-positive neurite. All cell lines showed a typical neuronal morphology and the presence of synaptic markers on neurites, indicating that despite their initially different differentiation speed both SHANK3-deficient and SHANK3-normal cells eventually differentiated into neurons (**Figure 2A-B**). Quantitative image analysis revealed several links to hyperdifferentiation in the SHANK3-deficient cell lines ASD04 and HT3. For example, we observed total MAP2 network length increases in SHANK3-deficient neurons (**Figure 2A-B**). Additionally, both SHANK3-deficient cell lines showed a significant increase of pre-synaptic Synapsin 1 spot numbers as well as a non-significant trend towards increased post-synaptic SHANK3 spots numbers (**Figure 2A-B**). Synapsin 1/SHANK3 double positive synapses were significantly increased in the HT3 neurons, while the increase in ASD04 neurons was less pronounced (**Figure 2A-B**). It is important to note that although SHANK3 spot numbers were increased, in line with our previous measurements of total cellular SHANK3 content by Western blotting and qPCR, also the SHANK3 signal intensity in Synapsin 1/SHANK3 synapses was decreased in SHANK3 deficient neurons (**Figure S1**, **Figure S2A**). Multi-Electrode Array (MEA) measurements further confirmed that all tested cell lines were electrophysiologically active, indicating their successful differentiation into neurons. However, in line with potentially structurally incomplete synapses, the electrophysiological activity was lower in the SHANK3-deficient cell lines ASD04 and HT3. Compared to the respective control cell lines at differentiation D35 the mean firing rate was reduced by approximately 50% and network oscillations were reduced by 30% (**Figure 2C**).

**Figure 2:**
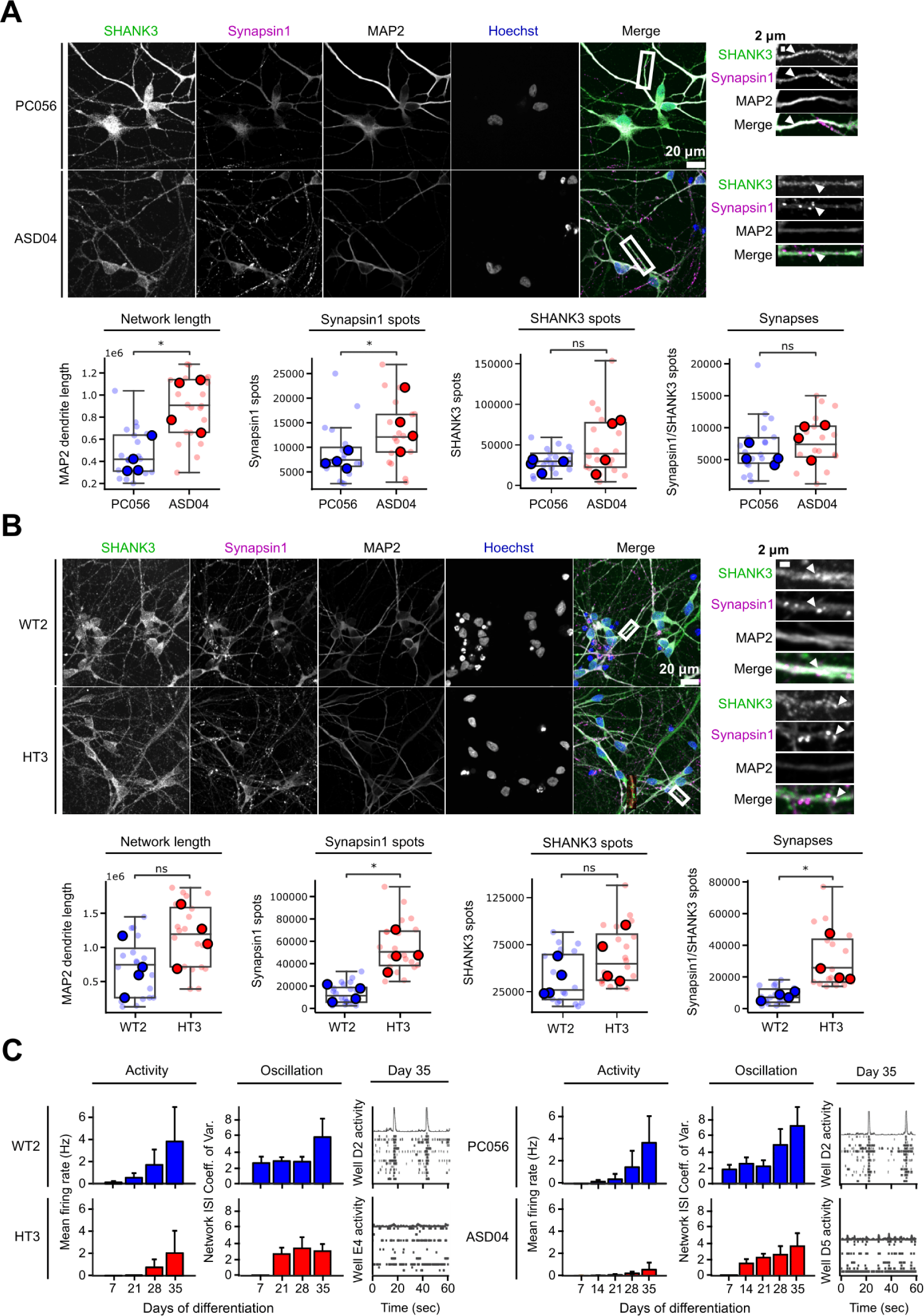
SHANK3-deficiency leads to increased neurite network length and number of synapses, but decreased neuronal activity. (A) The PMDS patient-derived ASD04 and healthy control PC056 NPC lines were differentiated for 30 days and fluorescently stained. The length of the MAP2 neurite network and the number of pre-synaptic Synapsin 1 spots, post-synaptic SHANK3 spots and Synapsin 1/SHANK3 double positive synapses were quantified. (B) Same as in (A), but in CRISPR-engineered SHANK3-deficient HT3 and isogenic control WT2 NPC lines. Each datapoint represents technical replicate data from one 384-well plate well. Larger markers represent the medians of independent differentiation experiments. The boxplots show the median and the 2^nd^ and 3^rd^ quartiles of the data. Whiskers indicate 1.5X of the interquartile range. Welch’s unequal variances t-test was performed using the medians of independent differentiation experiments. ns = not significant, * = p < 0.05. (C) Neuronal activity was measured at multiple differentiation timepoints in PMDS patient-derived and CRISPR-engineered cell lines using a Multi-Electrode Array (MEA). Activity represents the frequency of action potentials. Oscillations are alternating periods of high and low activity. Bar plots show data means from at least six MEA wells and error bars indicate the standard deviation. Raster plots show single well examples of neuronal activity over 60 seconds. Each row shows activity data from one electrode. Coordinated activity over multiple electrodes is expressed as a spike in the top row.

### Identification of hyperdifferentiation rescuing chemical compounds

Small molecules are frequently used to increase the efficiency of neuronal *in vitro* differentiation (23–25). We hypothesized that the observed SHANK3-specific increased differentiation during the early stages of cortical differentiation might also be amendable to modulation by small molecules. Accordingly, we designed a screening funnel to identify chemical compounds that decrease the observed hyperdifferentiation in a SHANK3-dependent manner in CRISPR-engineered and patient-derived NPCs (**Figure 3**). For primary screening, the CRISPR-engineered and SHANK3-deficient HT3 cell line was seeded at 3,500 cells per well in 384-well plates and treated with DMSO or 1 μM of 7,120 chemical compounds from D0 to D6. Cells were fixed and stained with Hoechst and antibodies against Ki67 and HuC/D. Images were segmented and the ratio of proliferative and differentiating cells was measured.

**Figure 3:**
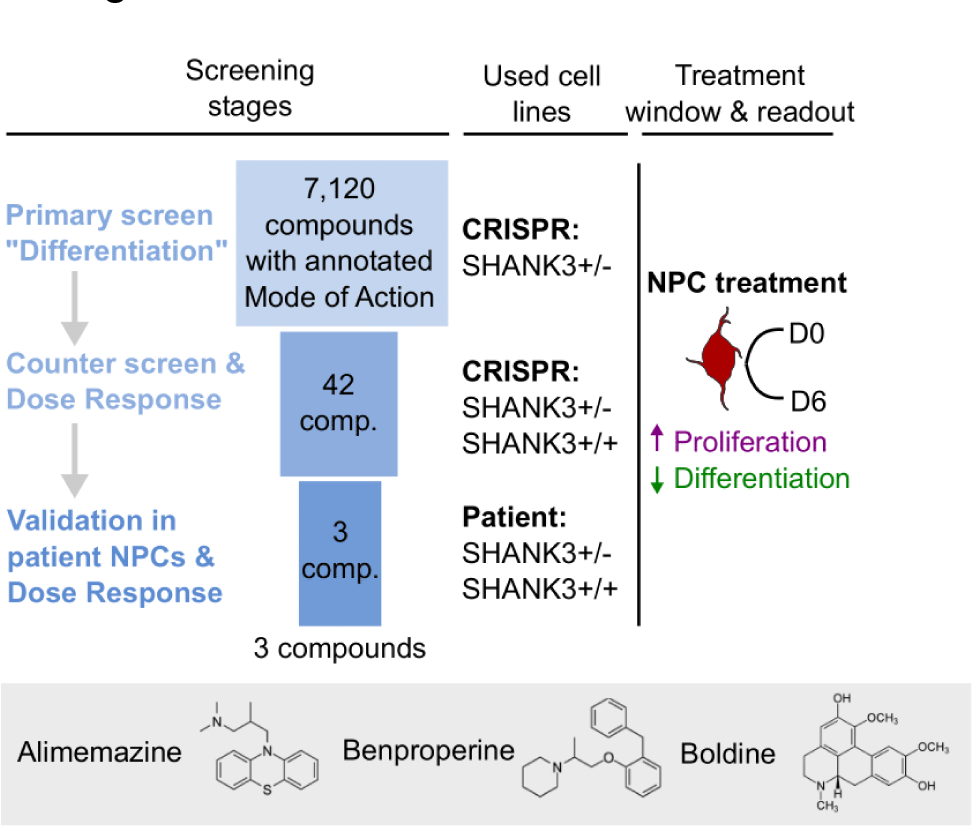
Overview of the applied screening strategy. During primary screening 7,120 Mode of Action (MoA)-annotated compounds were assessed based on non-toxic proliferation increase and differentiation decrease in the CRISPR-engineered HT3 SHANK3-deficient NPC line. Dose-response testing and counter screening in isogenic control WT2 and PMDS donor-derived NPCs yielded three candidate compounds.

22% of DMSO-treated NPCs were Ki67 positive and assumed to be proliferating, while 25% of NPCs were HuC/D positive and assumed to be differentiating (**Figure 4A**). We calculated a 3-standard deviation (SD) window around the DMSO control’s Ki67 and HuC/D ratio means. Compounds in the top left quadrant were considered as hits since those compounds increased proliferation and decreased differentiation compared to the DMSO control treatment. Compounds that fulfilled this criterium but resulted in a nuclei count < 2,000 were considered toxic and excluded. In total 42 compounds were identified that significantly increased proliferation and decreased differentiation (**Table S3**). The inhibitors HA-1077 and GSK-25 were used as positive controls. We measured average Z′ factors of 0.56 and 0.47 between GSK-25 and DMSO treated wells across all plates for the Ki67 and HuC/D ratio means, respectively (**Figure S3A-B**). Plates with a Z′ factor < 0.2 were discarded and repeated. The screen was performed in duplicate and was highly replicable as indicated by Pearson’s correlation coefficients between both replicates of 0.85 and 0.79 for the Ki67 and HuC/D positivity ratios, respectively (**Figure S3C**). Since we used a proliferation/differentiation readout, as expected, 21 out of 42 hits have annotated targets related to cell cycle regulation or differentiation such as mitogen-activated protein kinases (MAPKs), cyclin-dependent kinases (CDKs), rho-associated coiled-coil-containing protein kinase (ROCK), or glycogen synthase kinase-3 (GSK-3). Querying the STRING database with all identified targets showed a close physical or functional interaction for most (**Figure 4B**) (26). 5 detected hits function as GSK3-ß inhibitors and 4 hits as ROCK1/2 inhibitors. A number of target annotations occurred only once and were not connected to other annotated targets. Often those were related to neuronal signaling or remodeling such as acetylcholinesterase (AChE) inhibition, dopamine D_2_ receptor (DRD2) or histamine H_1_ receptor (HRH1) antagonism or actin-related protein 2/3 complex subunit 2 (ARPC2) inhibition. Next, we performed dose-response testing using all 42 identified hit candidates. We used the same CRISPR-engineered SHANK3-deficient cell line HT3 as before and additionally the isogenic control cell line WT2 expressing SHANK3 from both alleles to identify SHANK3 deficiency-specific molecules. Cells were treated from D0 to D6 in seven 3-fold dilution steps from 10 μM to 0.01 μM, fixed and stained with antibodies against Ki67 and HuC/D (**Figure 4C**). Compounds were selected based on three criteria: First, a dose-dependent increase in proliferating Ki67 positive cells, second a dose-dependent decrease in differentiating HuC/D positive cells, and third a SHANK3-deficiency specific effect. 11 compounds showed a SHANK3-defiency specific proliferation increase or were toxic in the SHANK3+/+ control cells (< 2,000 living cells) so that only mutant cell responses were evaluated (**Figure 4D**). Out of these 11 compounds, the three compounds Alimemazine, Benproperine, and Boldine showed also a SHANK3-deficiency specific effect on neuronal differentiation (**Figure 4E**). Out of the originally identified 42 compounds we therefore validated three compounds to have dose-dependent and SHANK3 deficiency-specific effects on NPC differentiation (**Table S1**).

**Figure 4:**
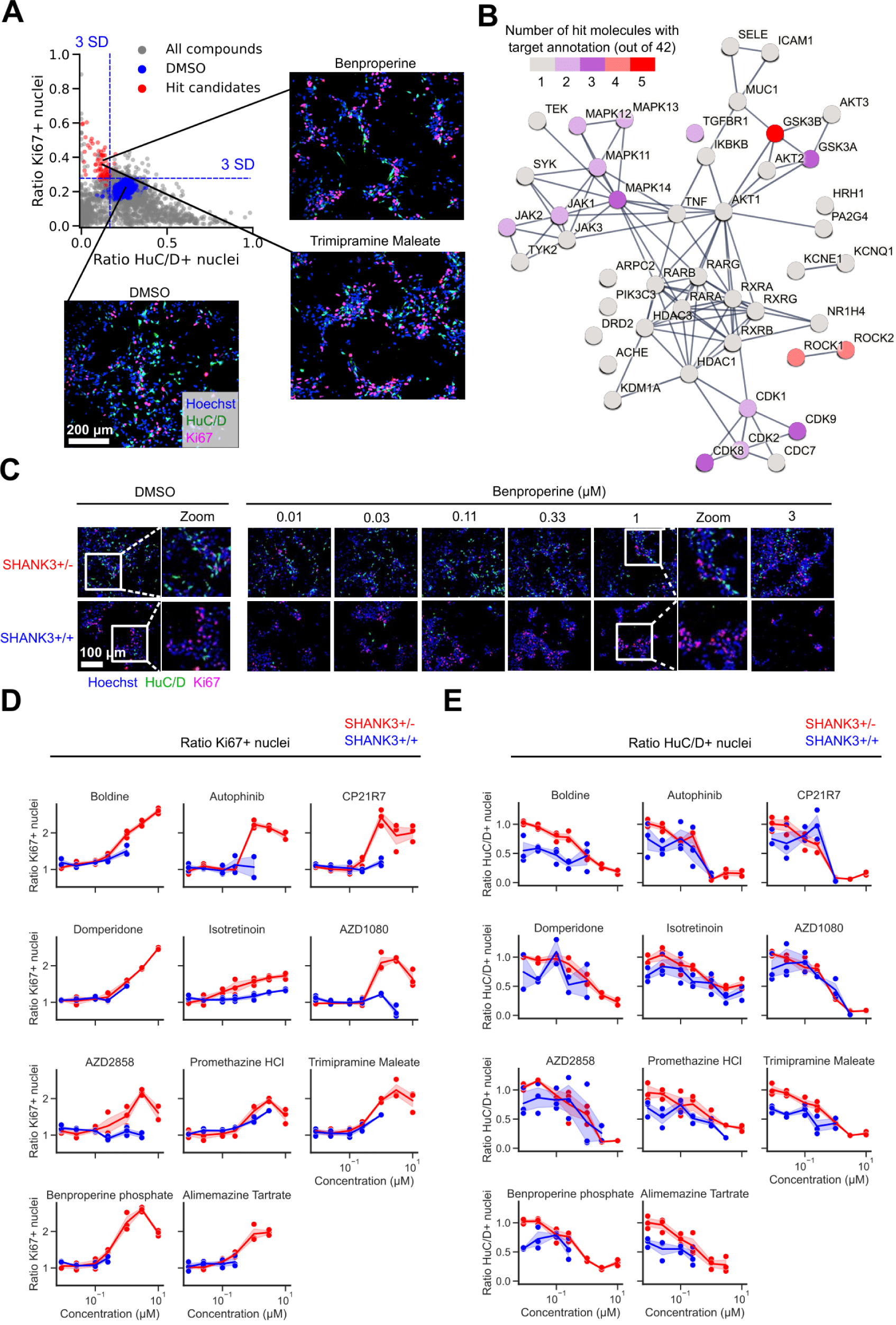
Identification of SHANK3 deficiency-specific hyperdifferentiation rescuing chemical compounds in CRISPR-engineered NPCs. (A) The top left quadrant shows 42 selected hit candidates (red) based on a 3 standard deviation (SD) window around the DMSO control’s Ki67 and HuC/D ratio means. Molecules that increased proliferation and decreased differentiation and did not lower the nuclei count < 2,000/well were considered as hit candidates. The primary screen was performed in duplicate and all data is shown. (B) Visualization of known physical and functional interactions of all protein targets attributed to the detected hit candidates annotated in the STRING biological database. The color coding represents the frequency of target annotations in the hit list. Compounds can have multiple targets (see also **Table S3**). (C) Representative images of CRISPR-engineered SHANK3+/- and control SHANK3+/+ NPCs treated with Benproperine at multiple doses. (D) Dose-response effects of 11 compounds on the ratio of proliferating Ki67 positive SHANK3+/- and control SHANK3+/+ NPCs. (E) Dose-response effects of 11 compounds on the ratio of differentiating HuC/D positive SHANK3+/- and control SHANK3+/+ NPCs. Concentrations that were toxic were not used for ratio calculations. The shaded area in the line graphs corresponds to the 95% confidence interval of triplicate experiments.

### Validation of SHANK3-specific hyperdifferentiation rescuing chemical compounds in patient-derived NPCs

Next, we validated the effects of the three identified compounds in the ASD04 PMDS patient-derived cell line (**Table S1**). The healthy donor derived PC056 cell line was used as control. Identical to previous experiments, cells were treated from D0 to D6 in seven 3-fold dilution steps from 10 μM to 0.01 μM, fixed and stained with Hoechst and antibodies against Ki67 and HuC/D (**Figure S4A**). The hyperproliferation and hypodifferentiation-inducing effects of all three tested compounds were confirmed in the SHANK3-deficient ASD04 NPCs. When comparing these results to the PC056 healthy-donor control NPC results, we could show that all three tested compounds specifically upregulated proliferation in a SHANK3 deficient manner (**Figure S4B**). SHANK3 deficiency-specific downregulated differentiation was also demonstrated in all three compounds (**Figure S4C**). For comparison, we also included three compounds (Domperidone, Promethazine, Trimipramine) that were not selected as SHANK3 deficiency-specific hits based on the previous results obtained in the CRISPR-engineered NPCs (**Figure 4D-E**). Although Domperidone, Promethazine, and Trimipramine showed SHANK3 deficiency-specific increases on NPC proliferation, we detected no clear SHANK3 deficiency-specific effects on differentiation. All three compounds decreased differentiation in both the ASD04 PMDS patient-derived cells as well as the healthy donor derived PC056 cells (**Figure S4B-C**). In conclusion, we identified Alimemazine, Benproperine and Boldine out of the 42 primary screening hits as chemical compounds that increase proliferation and decrease differentiation specifically in CRISPR-engineered as well as human PMDS patient-derived SHANK3 deficiency models (**Table S1**).

### Pharmacological hyperdifferentiation rescue depends on residual SHANK3 levels

PMDS patients are heterozygous SHANK3 mutation carriers, and the used SHANK3 mutated cell lines were designed to express approximately half of normal SHANK3 levels, but we also wanted to test whether the identified compounds require SHANK3 expression at all to show the identified proliferation and differentiation phenotypes (**Figure S1**). To examine the compounds’ SHANK3 dependence we used a CRISPR-engineered SHANK3 double knockout cell line (SHANK3-/-) and tested all three counter-screen validated compounds in these SHANK3-/-NPCs. Identical to previous experiments, cells were treated from D0 to D6 in seven 3-fold dilution steps from 10 μM to 0.01 μM, fixed and stained with Hoechst and antibodies against Ki67 and HuC/D. Compound toxicity was determined as increased nuclear staining intensity and decreased nuclear surface. SHANK3-/-double knockout cells were more susceptible to compound-induced toxicity than the previously tested cell lines (**Figure S5A**). Alimemazine and Benproperine lead to a decrease in nuclei number already starting at 0.37 μM, while Boldine showed decreasing cell counts at 1.12 μM. When only the non-toxic doses for each compound were considered, we saw no or only very small dose-dependent effects on proliferation or differentiation in the SHANK3-/-NPCs (**Figure S5A**). Boldine showed a small dose-response effect at non-toxic concentrations. Due to the absence of large compound-induced effects in the SHANK3-/-NPCs, we therefore conclude that residual SHANK3 levels are required for the full response to the identified compounds.

### Actin-related protein 2/3 complex subunit 2 (ARPC2) is expressed in D6 NPCs and its localization is altered by Benproperine treatment

Out of the three dose-response validated compounds only Benproperine was annotated with a protein target (**Table S1**). Benproperine was originally developed as a cough suppressant and is an actin-related protein 2/3 complex subunit 2 (ARPC2) inhibitor. Probably linked to actin-remodeling, Benproperine also suppresses cancer cell migration and tumor metastasis (27). To confirm whether ARPC2 is expressed in our NPC models, we performed TaqMan-probe RT-qPCR in D6 NPCs and measured the expression relative to the housekeeping gene *GAPDH*. Additionally we measured the mRNA expression levels of Dopamine receptor D2 (*DRD2*) and Histamine Receptor H1 (*HRH1*), the annotated targets of the compounds Domperidone and Promethazine which were not selected as hits (**Figure 5A**). In both WT2 and HT3 cell lines we found that *DRD2* and *HRH1* were expressed at 1% or less of the *GAPDH* mRNA level. In contrast, *ARPC2* was expressed between 10-31% of the *GAPDH* mRNA level. Furthermore, RT-qPCR showed that ARPC2 inhibitor Benproperine did not impact the *ARPC2* mRNA expression level (**Figure S5B**). ARPC2 is a subunit of the seven-subunit Actin Related Protein 2/3 (ARP2/3) complex involved in the regulation of the actin cytoskeleton and specifically neurite outgrowth and corticogenesis (28,29). Since no other ARP2/3 complex inhibitors were part of the 7,120 screened molecules, we tested CK-666, a classic ARP2/3 complex inhibitor, to show that Benproperine exerts its proliferation inducing effect via the ARP2/3 complex (30). As before we measured the ratio of Ki67 positive NPCs as a proxy for proliferation. Similar to our previous results, we found that at D6 HT3 NPCs proliferated less than WT2 NPCs and that this phenotype was rescued by a 6-day treatment with 1 μM Benproperine. Although less than Benproperine, also CK-666 lead to a dose-dependent increase of HT3 NPC proliferation suggesting a possible link to ARP2/3 complex inhibition (**Figure 5B**). Since the ARP2/3 complex is closely physically associated with actin, we next investigated whether Benproperine would have an impact on ARPC2 localization or expression on the protein level. We performed immunofluorescence experiments in D6 NPCs using antibodies against ARPC2, ß-actin, and ß-tubulin III (**Figure 5C**). We observed ß-actin accumulation in ß-tubulin III-positive neurites and colocalization with the ARPC2 signal. Using colocalization analysis we found that ARPC2/ß-actin colocalization (32%) was lower in the SHANK3-deficient HT3 cell line (20%) (**Figure 5D**). Treatment with Benproperine rescued the lowered colocalization to 30%. It is important to note that this effects is also SHANK3-deficiency specific and was only observed in the HT3 NPCs. The protein levels of ARPC2 and ß-actin, as measured by the mean fluorescent intensity in the images, were not different between the WT2 and HT3 cells and Benproperine treatment did not change ARPC2 or ß-actin protein levels in line with our qPCR results (**Figure 5D**, **Figure S5B**). Benproperine therefore impacts ARPC2/ß-actin colocalization without altering protein expression.

**Figure 5:**
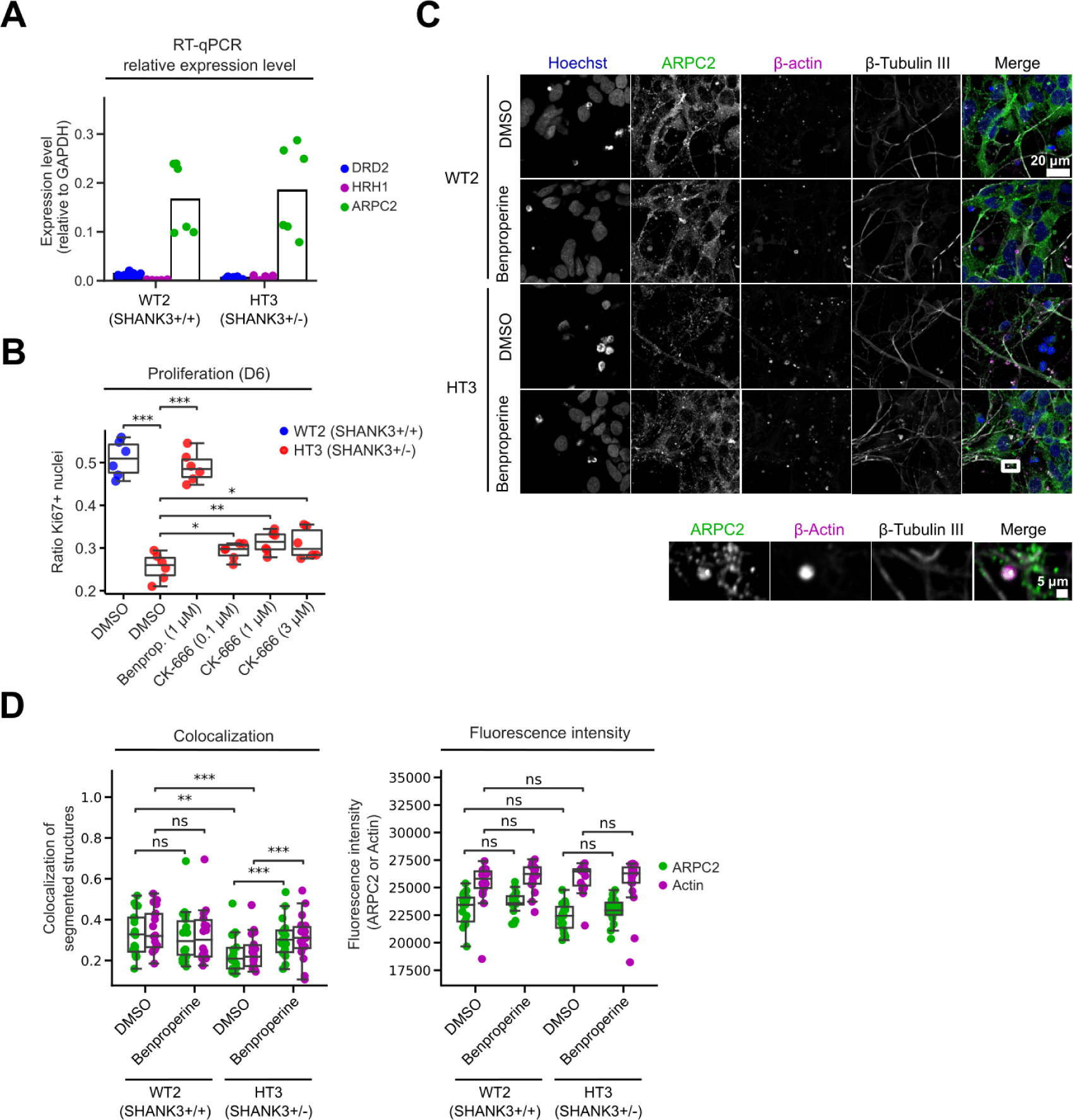
Actin-related protein 2/3 complex subunit 2 (ARPC2) inhibition increases proliferation and alters ARPC2 colocalization with ß-actin. (A) RT-qPCR was performed using Taqman probes against the annotated compound targets dopamine-2 receptor (*DRD2*), histamine H1 receptor (*HRH1*) and actin-related protein 2/3 complex subunit 2 (*ARPC2*). The expression level was determined relative to *GAPDH*. (B) Measurement of Ki67 positive proliferating WT2 and HT3 NPCs treated with the ARPC2 inhibitors Benproperine and CK-666. (C) Representative images of DMSO or Benproperine treated D6 NPCs with ARPC2, ß-actin, and ß-tubulin III immunofluorescent staining. (D) Quantification of ARPC2 and ß-actin colocalization and fluorescent intensity. Two independent differentiations were performed using multiple technical replicates. The boxplots show the median and the 2^nd^ and 3^rd^ quartiles of the data. Whiskers indicate 1.5X of the interquartile range. Welch’s unequal variances t-test was applied. ns = not significant, * = p < 0.05, ** = p < 0.01, *** = p < 0.001.

### The ARPC2-targetting compound Benproperine rescues increased synapse numbers when administered early during differentiation

Given the role of the ARP2/3 complex for neurite outgrowth via actin remodeling in neuronal growth cones, we investigated Benproperine effects on synapse formation and neuronal activity. We treated CRISPR-engineered WT2 and HT3 cells with 1 μM Benproperine or DMSO for 72 hours at either one of two different timepoints during neuronal differentiation. An early treatment interval from D1-D3 was chosen to evaluate Benproperine’s effect during development on synapse formation and activity, while a later interval from D34-36 in previously untreated neurons was chosen to test Benproperine’s effects on already formed synapses. Neurons were stained with Hoechst, and antibodies against pre-synaptic marker Synapsin 1, post-synaptic protein SHANK3 and MAP2 (**Figure 6A**). Consistent with our previous results, we measured an approximately two-fold increased number of synapses in HT3 neurons compared to WT2 control neurons (**Figure 2A-B**, **Figure 6B**). Interestingly, only an early treatment from D1-3 lowered the synapse count in HT3 neurons, thereby partially rescuing increased synapse formation. Later treatment from D34-36 did not lower synapse counts compared to DMSO. In line with our previous findings, Benproperine therefore decreased neuronal differentiation leading to reduced synapse formation while not being synaptotoxic when administered later during differentiation. We also tested the effects of the other two identified hits Alimemazine and Boldine on synapse formation and observed that both compounds impacted synapse numbers in HT3 neurons. While Alimemazine increased synapse formation, Boldine decreased the number of synapses similar to Benproperine (**Figure S6A**). To understand whether the rescue of synapse numbers has also effects on neuronal activities we differentiated WT2 and HT3 neurons in MEA plates for 42 days and measured their electrical activity in regular intervals. In line with our previous results we found that untreated WT2 neurons were more active than SHANK3-deficient HT3 neurons on the individual neuron level, but also on the network level (**Figure 2C**, **Figure 6C-D**). While the mean neuronal firing rate of WT2 neurons was about 10 Hz, it was reduced to 3 Hz in HT3 neurons. Also signal synchrony, reflecting the strength of synaptic or inter-electrode connections, and network oscillation, reflecting the coordination of network activity across electrodes, were decreased in HT3 neurons (**Figure 6D**). When Benproperine was added either early (D1-3) or late (D40-42) during WT2 and HT3 neuron differentiation, interestingly only minor changes were observed. In HT3 neurons the neural activity, as measured by firing rate, was slightly decreased from 3 to 2 Hz when Benproperine was administered early during differentiation (D1-3), but not during late treatment (D40-42). Higher order neuronal synchrony and network oscillation patterns remained unchanged after Benproperine treatment (**Figure 6D**). SHANK3 is a crucial scaffolding protein in the postsynaptic density and we hypothesized that a compound-induced lack of SHANK3 might be associated with the observed reduction of synapses and the mild decrease of neuronal firing rate. As described above, we first determined synapses as Synapsin 1/SHANK3 double positive spots on the MAP2 network in D36 neurons. As an indicator of the SHANK3 protein level in synapses the integrated sum of all SHANK3 channel pixel intensities specifically within synapses was determined. As measured before, we observed that the synapses in untreated SHANK3-deficient HT3 neurons contained less SHANK3 signal (**Figure S2A**, **Figure 6E**). When we measured the synaptic SHANK3 signal in Benproperine-treated neurons, to our surprise, the SHANK3 integrated intensity was lower in both tested cell lines and there was no difference between both cell lines (**Figure 6E**). Benproperine therefore seems to affect synaptic SHANK3 levels less in SHANK3-deficent HT3 neurons.

**Figure 6:**
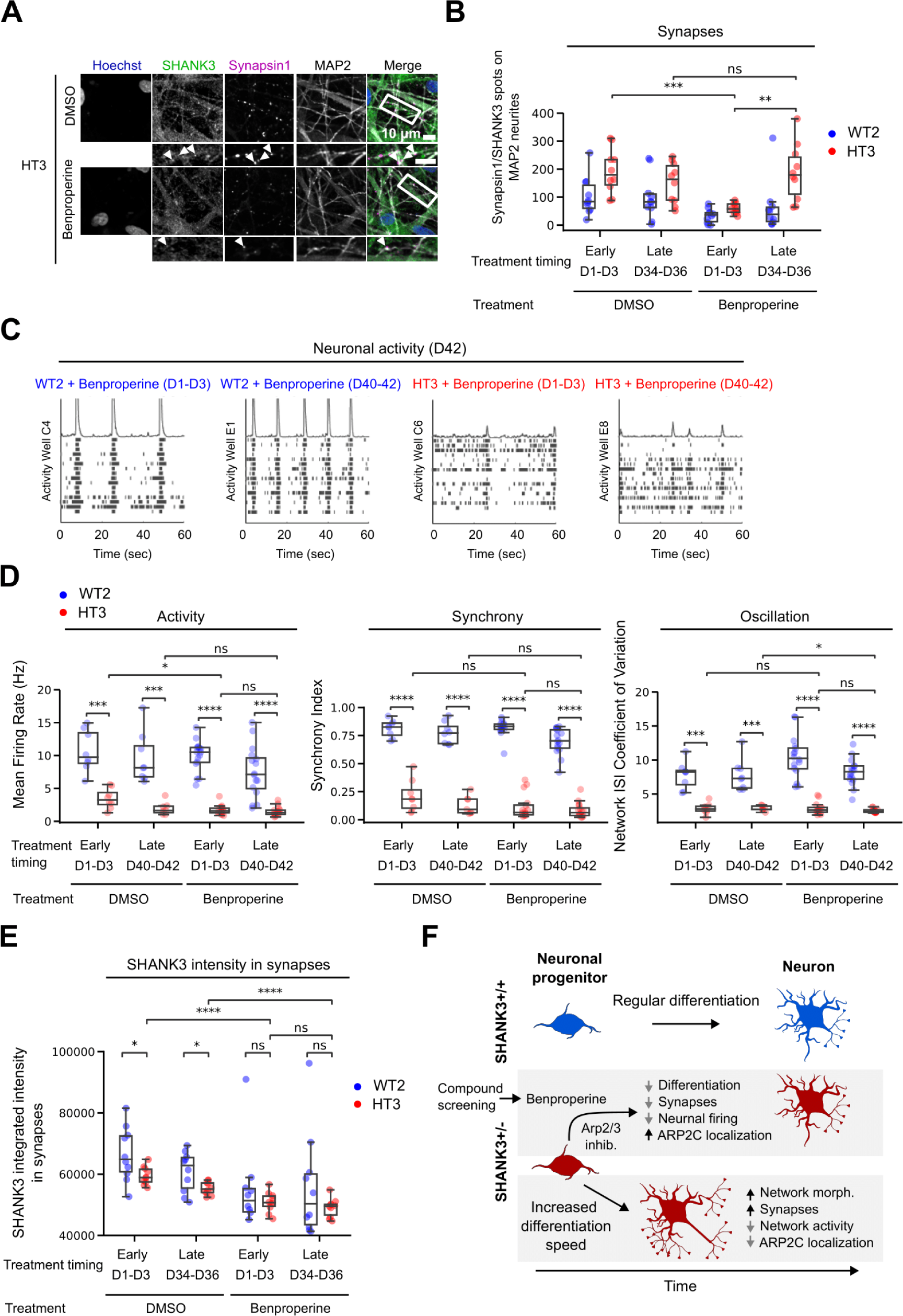
Benproperine modulates synapse formation and neuronal activity. (A) Representative images of fluorescently stained CRISPR-engineered SHANK3-deficient HT3 neurons after 36 days of differentiation and Benproperine treatment. (B) HT3 mutant or WT2 isogenic control cells were treated with Benproperine either early (D1-3) or late (D34-36) during differentiation and synapses were quantified. (C) Representative raster plots showing HT3 mutant or WT2 isogenic control neuron activity on a Multi-Electrode Array (MEA) plate after 42 days of differentiation. The cells were treated with Benproperine either early (D1-3) or late (D40-42) during differentiation. (D) WT2 and HT3 neuronal activity, synchrony, and oscillation determined by MEA after 42 days of differentiation and after early (D1-3) or late (D40-42) Benproperine treatment. (E) Quantification of the integrated SHANK3 pixel intensities in synapses (Synapsin 1/SHANK3 double positive puncta on MAP2 dendrites). (F) Schematic overview of the effects of SHANK3 haploinsufficiency and Benproperine treatment on neuronal differentiation. Two separate differentiations were performed. The boxplots show the median and the 2^nd^ and 3^rd^ quartiles of the data. Whiskers indicate 1.5X of the interquartile range. Welch’s unequal variances t-test was performed. ns = not significant, * = p < 0.05, ** = p < 0.01, *** = p < 0.001.

Taken together, our results highlight that Benproperine might act via an inhibition of the ARP2/3 complex mainly early during differentiation by increasing NPC proliferation, which inhibits synapse formation due to a developmental delay. Additionally, Benproperine can impact synaptic SHANK3 content. In line with these findings also individual neuronal activity is decreased, while the overall neuronal network is still functional. Benproperine administration to mature neurons has no effects on synapse numbers or neuronal activity, highlighting the compound’s specific activity during neurodevelopment (**Figure 6F**).

## Discussion

In this study we used PMDS patient and CRISPR-engineered human neuronal PMDS models and show that SHANK3-deficiency leads to decreased proliferation and increased differentiation of human NPCs (**Figure 1**). Increased differentiation also increased neuronal network length and the number of synapses, while decreasing neuronal activity (**Figure 2**). In various model systems and different SHANK3 genetic backgrounds others have described related results. In Shank3B^−/−^ knockout mice, Peixoto *et al.* also found earlier maturation of spiny projection neurons and increased corticostriatal network activity (9). Using human ASD patient derived neurons carrying SHANK3 microdeletions, an additional study showed that the number of primary neurites was increased when SHANK3 levels were decreased (8). Another group also found a decrease of proliferating cells in human SHANK3-deficient NPCs (7). Given its classical localization to the PSD, the question remains how SHANK3 is involved in the regulation of differentiation before the PSD has formed. We performed a screen using 7,120 molecules with annotated MoAs to identify small molecular compounds that can reverse the observed hyperdifferentiation (**Figure 3**). 21 out of 42 obtained hits had annotated targets related to cell cycle regulation or differentiation such as MAPKs, CDKs, ROCK, or GSK-3 (**Figure 4B**). This target enrichment is most likely due to the used Ki67 and HuC/D proliferation and differentiation readout. When we performed counter screening in both healthy donor and non-modified CRISPR SHANK3 control NPCs none of these 21 cell cycle regulation and differentiation-related molecules showed a SHANK3 deficiency specificity and were discarded (**Figure S4**, **Table S1**). After these selection steps the three molecules Alimemazine, Benproperine, and Boldine were found to rescue hyperdifferentiation without toxicity in a SHANK3 deficiency-specific manner.

Since actin remodeling is of relevance to neuronal differentiation we focused on the ARP2/3 complex subunit ARPC2 inhibitor Benproperine. Benproperine rescued ARPC2 colocalization to β-actin in developing dendrites while not changing total protein levels (**Figure 5C-D**). Several known genetic risk factors for psychiatric disease are related to the regulation of actin polymerization (29). Although it is unclear how SHANK3 deficiency influences cortical development, SHANK3 interactors, such as ARP2/3 complex subunits, that modulate actin polymerization have been described to impact cortical development and synaptic plasticity. Differentiation-linked processes such as morphological changes or lineage specification depend on the activity of the ARP2/3 complex (31,32). Next to increased neurite formation, Kathuria *et al*. found reduced actin levels in SHANK3-deficient human neurons (8). Direct SHANK3 interactions with the ARP2/3 complex and actin in transgenic mice, human cell lines, and Zebrafish have been shown to modulate actin levels and to influence dendritic spine development (33–35). Additionally, the structural plasticity of dendritic spines is negatively impacted by the deletion of ARPC2/3 complex subunit ARPC3 with consequences for cognition and social behavior in mice (36). Also other actin regulators can cause SHANK3-related autism phenotypes. For example, the aberrant regulation of synaptic actin filaments by the actin-binding protein cofilin has been shown to contribute to the manifestation of autism-like phenotypes in mice, while modulation of the cofilin upstream kinases PAK and Rac1 partially rescued social deficits (33). Taken together, it is therefore plausible that a chemical compound which targets actin polymerization via ARPC2 inhibition can rescue neuronal hyperdifferentiation, thereby normalizing synaptic density on dendrites. Supporting a close connection between SHANK3, the ARP2/3 complex and differentiation is our finding that Benproperine requires residual levels of SHANK3 protein and fails to rescue hyperdifferentiation in full SHANK3 knockout cells (**Figure S5A**).

We found that the SHANK3 protein level in synapses is lower in SHANK3+/-cells (**Figure S2A**, **Figure 6E**). In line with our findings mRNA imaging previously showed that *SHANK3* mRNA content is lower in the processes of neurons derived from a SHANK3 haploinsufficient donor (37). Additionally, it is important to note that Benproperine did not rescue SHANK3 synaptic protein levels, but rather lead to a further decrease. The decrease was, however, more pronounced in WT2 neurons than in HT3 neurons resulting in an equal synaptic SHANK3 content after treatment early or late during differentiation. Considering the important scaffolding function of SHANK3, the observed compound-induced decrease of synapse formation and mild decrease of neuronal firing could also be the consequence of a disturbed synapse structural integrity due to further reduced SHANK3 levels (**Figure 6B-D**). Most of the described Benproperine-associated phenotypes were milder in SHANK3+/+ cells. This might be due to the overall higher SHANK3 protein level in these cells which buffers Benproperine effects. Additionally, future research needs to address whether the ARP2/3 complex or actin remodeling is involved in synaptic SHANK3 level maintenance, for example due to mRNA transport.

Benproperine only decreased Synapsin 1/SHANK3 synapses when NPCs were treated early (D1-3), but not late during differentiation (D34-36) (**Figure 6B**). MEA measurements showed that Benproperine only mildly affected neuronal firing and in line with the synaptic data only when administered early during differentiation (**Figure 6D**). Other chemical compounds have also been described to have differentiation blocking effects in NPCs. For example, several antidepressants have been shown to stimulate adult NPC proliferation (38–40). Similar to Benproperine, the proliferation stimulating effect of Lithium depends on the genetic background of the patient (41). For obvious reasons, any future PMDS treatment strategy targeting neuronal differentiation will face high safety hurdles and will require careful compound characterization. We anticipate that human PMDS stem cell models will play a key role in developing novel molecular treatment hypotheses and to identify and characterize novel chemical neurodevelopmental modulators.

## Methods

### Neuronal progenitor cell generation

All cell lines that were used in this study are described in **Table S4**. PC56 iPSC line was obtained commercially at Phenocell (France). 4603 fibroblasts were obtained from Coriell’s biorepository (Coriell Institute for Medical Research, NJ, USA), PDF01 fibroblasts were obtained from ATCC repository (American Type Culture Collection, VI, USA). SHANK3-iPSC lines were derived from fibroblasts of 3 autistic children with de novo SHANK-3 mutations. The 3 patients were diagnosed with autism and severe intellectual according to DSM-IVTR criteria. They were included initially in a national observational study (IDRCB 2008-A00019-46) and, in this context, were submitted to whole genome sequencing to exclude additional autism-linked mutations. After patient’s legal representatives approval, 8-mm skin punch biopsies were obtained (study approval by Comity for the Protection of Persons CPP no. C07-33) and fibroblasts were isolated. Fibroblasts were reprogrammed using the four human genes OCT4, SOX2, c-Myc, and KLF4 cloned in Sendai viruses. The obtained iPSC lines were characterized as previously described (42). Commitment of the iPSCs to the neural lineage and derivation of stable late cortical progenitors (LCP) was described previously (23). Briefly, neural commitment was induced using the BMP inhibitor Noggin and the Nodal inhibitor SB431542. At day 10, neural rosettes containing neuro-epithelial cells were collected and plated en bloc in poly-L-ornithin/laminin treated culture dishes in N2B27 medium containing Epidermal Growth Factor (EGF), FGF-2, and Brain-derived Growth Factor (BDNF). At confluence, the passages were performed using trypsin at a density of 50,000 cells/cm². Mass amplification was performed until passage 8 and cells were frozen. To start the terminal differentiation as post-mitotic neurons, cortical NPCs were dissociated and plated in N2B27 without growth factors.

### Gene editing

SHANK3 gene–knockout mutants were generated using CRISPR/Cas9-mediated genome editing. Guide RNAs (sgRNAs) were designed using Crispor software (http://www.crispor.tefor.net). CRISPR/Cas9 was complexed with sgRNA and TracrRNA and inserted in the hESC line SA001 by nucleofection (**Table S4**). After nucleofection, cells were plated in Petri dishes containing StemMACS medium supplemented with Clone-R Supplement. After 1 week, cells were dissociated and sorted using the C.SIGHT clonal dispenser (Cytena) or the MoFlo cell sorter (Beckman) in 96-well plates. CRISPR/Cas9 editing was first screened using PCR and occurrence of deletions further validated by Sanger sequencing (Genewiz Facilty, Azenta Life Science). Edited hESC were further differentiated as described for iPSC.

### Cellular differentiation

Cryopreserved NPCs were thawed in basal medium (**Table S2**) and centrifuged at 300 g for 5 minutes. Cell pellets were resuspended in proliferation medium (**Table S2**) supplemented with ROCK inhibitor. Cells were seeded in T175 flasks coated with Poly-Ornithine/Laminin (**Table S2**) and incubated at 37°C and 5% CO_2_. 24 hours after thawing, the proliferation medium was changed to remove Rock inhibitor. After 3 days in proliferation medium, cells were detached, counted using an automated cell counter (Countess, ThermoFisher) and centrifuged at 300 g during 5 minutes. The cell pellet was resuspended in differentiation medium (**Table S2**). 384 well plates (Greiner, 781091) were coated with Poly-Ornithine using a Multidrop Combi (ThermoFisher) for dispensing and a Bravo automated liquid handler (Agilent) for PBS washes and Laminin dispensing. Both coatings were incubated overnight at 37°C and 5% CO_2_. Just before cell seeding, the coating solution was aspirated using a Bravo automated liquid handler leaving 10 µL of coating solution inside the well to ensure coating intactness. *Developmental phenotype:* After resuspension in differentiation medium, cell seeding was performed using a Multidrop Combi (ThermoFisher) adding 40 µL of cell suspension per well. Cells were incubated at 37°C and 5% CO_2_. In case of compound treatment, cells were treated 5 hours after plating by adding 10 µL of differentiation medium. 0.1% DMSO was used as negative control and 10 µM HA-1077 or 1 µM GSK-25 (**Table S2**) were used as positive controls for SHANK3+/+ or SHANK3+/-cells, respectively. On day 4 of differentiation, half of the medium (30 µL) was changed. Compound treatments were performed using a VPrep liquid handler (Agilent). Cells were fixed on day 6 and stained with Hoechst, and antibodies against Ki67 and HuC/D. *Synaptic phenotype:* After cell resuspension in differentiation medium, seeding was performed using a Multidrop Combi (ThermoFisher) in the following condition: 50 µL/well containing 500 SHANK3 +/+ cells/well and 2500 SHANK3 +/-cells/well. Cells were then incubated at 37°C, 5% CO2. Twice a week, medium was removed and 30 µL/well of fresh differentiation medium were added. From day 14, the differentiation medium was replaced by a BrainPhys medium (**Table S2**). On days 23 and 25, cells were treated with chemical compounds and 0.1% DMSO as vehicle control (**Table S2**). On day 28, cells were fixed and stained with antibodies against MAP2, Synapsin 1 and SHANK3 expression.

### RT-qPCR

Cells were seeded and differentiated as described before until D6 or D28. qPCR samples were prepared using the Fast Advanced Cells-to-CT TaqMan kit (ThermoFisher) according to the manufacturer’s protocol. Expression levels were determined by relative quantification using the ΔΔCt method. *SHANK3* mRNA relative expression was determined in comparison to *PPIA*, described before as a suitable control (17). Expression levels of *DRD2*, *HRH1*, and *ARPC2* were determined relative to *GAPDH* and *PPIA* expression.

### Western blotting

Cell lysis was performed in RIPA buffer. Samples were loaded in NuPAGE LDS Sample Buffer and ran on NUPAGE Novex 3-8% Tris-acetate gels immersed in NUPAGE Tris-acetate SDS running buffer (1x) for 1.5 hours at 120V (**Table S2**). Gels were transferred to PVDF membranes using the Trans-Blot Turbo Transfer System (Biorad) and the high molecular weight transfer option. Membrane blocking was performed in 5% skim milk powder in TBS-T for 1 hour at room temperature. Used primary and secondary antibodies are summarized in **Table S2**. The anti-SHANK3 antibody was preincubated with a SHANK3 blocking peptide to demonstrate signal specificity. The membrane was developed using SuperSignal West Dura chemiluminescent substrate and imaged using an iBright (ThermoFisher) imager. Band intensity was determined using the Area Under the Curve (AUC) method in FIJI/ImageJ (43).

### Immunofluorescence

All steps were performed using an EL406 liquid handler (Biotek). Fixation was performed for 20 minutes by directly adding 20 µL/well of 16% PFA (4% final) to the cells, followed by three PBS washes. Blocking and permeabilization was performed for 1 hour in 1% BSA and 0.05% Triton X-100 diluted in PBS (**Table S2**). After a PBS wash, primary antibodies were added in PBS containing 1% BSA and incubated overnight at 4°C. After three PBS washes, the secondary antibodies were added in PBS containing 1% BSA and incubated for 90 minutes at RT, followed by three PBS washes (**Table S2**).

### Imaging and image analysis

Imaging was performed on a Yokogawa CV7000 microscope in scanning confocal mode using a dual Nipkow disk. 384-well plates were mounted on a motorized stage and images were acquired in a row-wise “zig-zag” fashion. The system’s CellVoyager software and 405/488/561/640nm solid laser lines were used to acquire 16-bit TIFF images through a dry 20X or water-immersion 60X objective lens using a cooled sCMOS camera with 2560×2160 pixels and a pixel size of 6.5μm without pixel binning. Using the 20X objective 9 single Z-plane images were acquired in a 3×3 orientation from the center of each well. Using the 60X objective 16 images from three 1 µm-separated Z-planes were acquired in a 4×4 orientation from the upper right corner of the well. Maximum intensity projection was performed. Image segmentation and feature extraction was performed with our in-house software PhenoLink (https://github.com/Ksilink/PhenoLink). Image segmentation was performed on illumination corrected raw images based on fluorescent channel intensity thresholds empirically determined per plate. Nuclei were segmented using intensity and size thresholds allowing the identification and counting of debris, dead cells and live cells. The ratio of HuC/D and Ki67 positive cells was calculated based on an intensity threshold for segmentation and a second colocalization threshold with the segmented nuclei (50% and 5%, respectively). Synapses were identified in a three step process: First a MAP2-channel based neurite skeleton was determined. Secondly, puncta fitting based on local maxima and individual puncta intensity profile were used to identify puncta in the Synpasin1 and SHANK3 channels. Puncta selection was performed using signal over noise, size and intensity thresholds. Lastly, synapses were identified by determining the triple colocalization of both synaptic puncta signals with the neurite skeleton.

### Screening and hit detection

The CRISPR-engineered and SHANK3-deficient HT3 cell line was seeded at 3,500 cells per well in 384-well plates as described above and treated in duplicate with 1 μM of 7,120 chemical compounds from Selleckchem’s Bioactive compound library 5 hours post-seeding from D0 to D6. Cells were fixed, stained, imaged and the ratios of proliferating and differentiating cells were calculated as described above. Data across screening runs was normalized per plate using median normalization based on the DMSO control wells. The Z′ factor between GSK-25 and DMSO treated control wells was calculated to assess plate quality. Plates with Z′ factor <0.2 were discarded. Hit candidates were defined as molecules that increased the ratio of proliferating and decreased the ratio of differentiating cells more than 3 standard deviations (SDs) from the median of the DMSO control. Compounds that lowered the live cell count below 2,000 per well were discarded as toxic. All raw data and the corresponding analysis workflow is available as a Jupyter notebook on GitHub (https://github.com/Ksilink/Notebooks/tree/main/Neuro/SHANK3screening). Potential biological interactions between annotated hit targets were explored and visualized using the STRING biological database. All available annotated target-related gene name inputs of all 42 primary hits are summarized in **Table S3**.

### Neuronal activity

Cryopreserved NPCs were thawed in a water bath and centrifuged (400g, 5 min, RT) in basal medium supplemented with ROCK inhibitor (**Table S2**). Cell pellets were resuspended in differentiation medium (Table S4) supplemented with ROCK inhibitor. 48-well CytoView MEA plates were coated with Polyethyleneimine solution (**Table S2**) for 1 hour at 37°C followed by Laminin coating for 2 hours at 37°C. 150×10^3^ cells/well were seeded in 15 µL in the center of each well on the electrodes and incubated for 1 hour at 37°C. After 1 hour, 300 µL of medium was added. Cells were incubated at 37°C and 5% CO2 for up to 42 days with differentiation medium changes twice a week. Before media changes, the electrical activity was recorded on the Maestro Pro Multiwell MEA (Axion Biosystems) using the AxIS Navigator software (version 3.5.1, Axion Biosystems). Electrical activity was recorded for 5 minutes at a 12.5 kHz sampling rate and a 5.5 standard deviation threshold level for action potential detection. Before plate loading the device was allowed to equilibrate for approximately 30 minutes to 37°C and 5% CO2. Data analyses were performed using the Neural Metric Tool (version 3.1.7, Axion Biosystems). We focused on three activity, synchrony, and oscillation, each calculated per well. Activity is the mean firing rate across electrodes, representing neuron function. Synchrony is the normalized area under the inter-electrode cross-correlation, reflecting synaptic connection strength. Oscillation is the coefficient of variation in inter-spike intervals, measuring whole network activity. High values indicate bursts of action potentials, while low values indicate uncoordinated action potentials across neurons.

## Author contributions

AT designed and performed differentiation and synaptic imaging experiments, performed primary screening, and analyzed imaging data; KS optimized the differentiation protocol, performed MEA analysis, and designed experiments; DH performed synaptic imaging, MEA experiments, and qPCR; LCo designed the image analysis workflow and analyzed imaging data; AW performed primary screening; AV performed synaptic imaging experiments; LF and LCh created cell lines; PS co-initiated the project; CJ supervised screening experiments; AB co-initiated the project; JW designed experiments, analyzed data, supervised the project, and wrote the manuscript with input from all authors.

## Acknowledgements

We thank Arnaud Ogier for help with screening data management and all colleagues at Ksilink for regular and helpful discussions. Ksilink is supported by the “Programme d’investissements d’avenir” (PIA) of the French government.

## Supplementary Figures

**Figure S1 referring to Figure 1:**
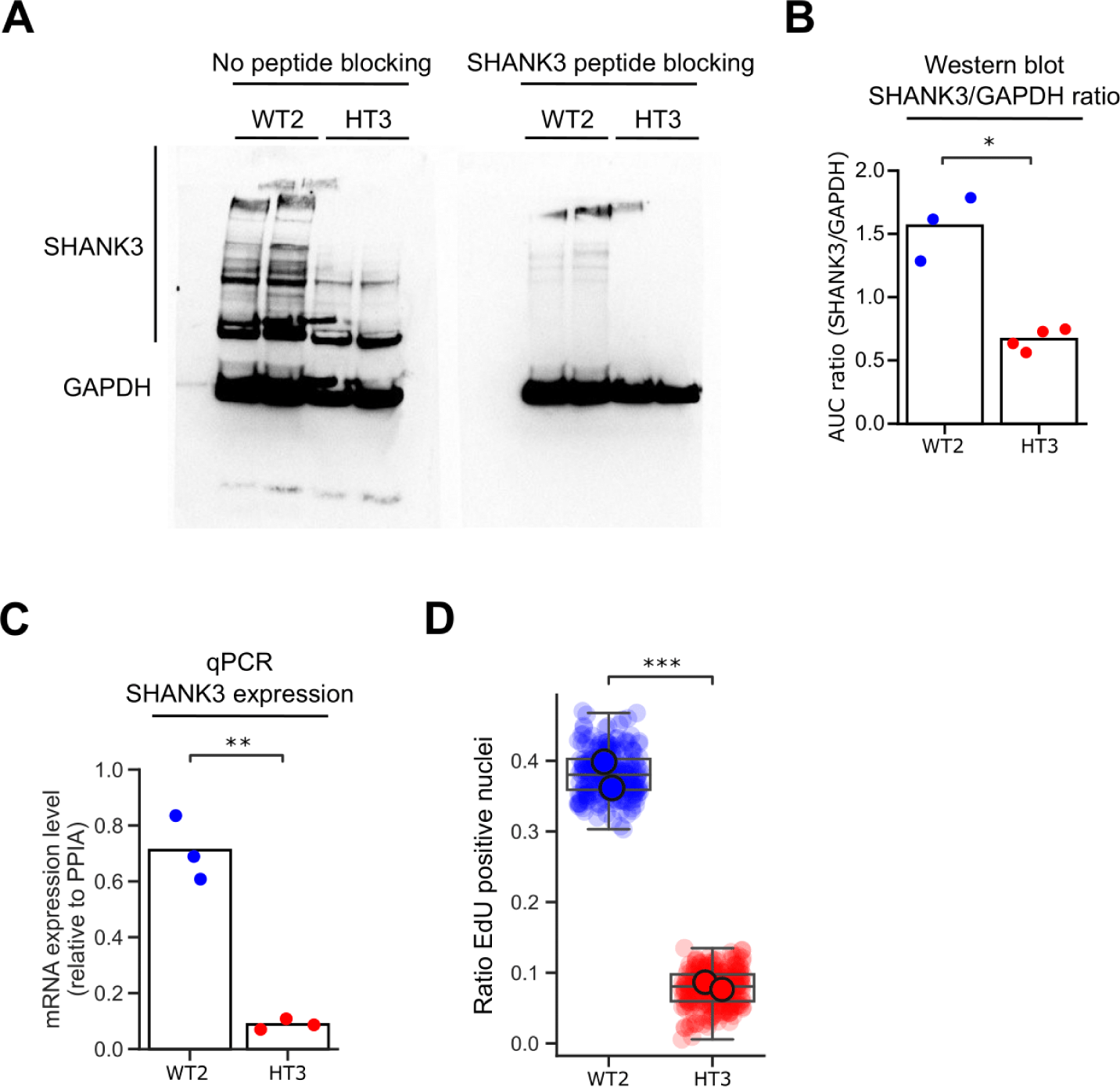
SHANK3 expression and proliferation is reduced in CRISPR-modified NPCs. (A) Representative images of Western blots with and without SHANK3 peptide blocking of WT2 and HT3 NPCs lysed around D30. (B) Quantification of the Western blot SHANK3 signal relative to the GAPDH signal using the Area Under the Curve (AUC) method in FIJI/ImageJ. (C) RT-qPCR was performed using Taqman probes against SHANK3 on D30. The expression level was determined relative to PPIA. At least two independent experiments were performed for Western blot and RT-qPCR experiments. (D) EdU (5-ethynyl-2’-deoxyuridine) incorporation assay in WT2 and HT3 cells at D14. Larger markers represent the medians of two independent differentiation experiments, smaller markers the technical replicates. The boxplots show the median and the 2^nd^ and 3^rd^ quartiles of the data. Whiskers indicate 1.5X of the interquartile range. Welch’s unequal variances t-test was performed using the medians of independent differentiation experiments. ns = not significant, * = p < 0.05, ** = p < 0.01, *** = p < 0.001.

**Figure S2 referring to Figure 2:**
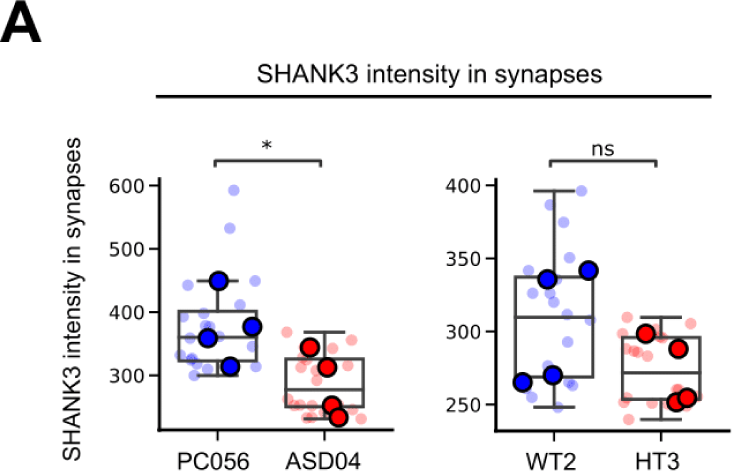
Synaptic SHANK3 levels are lower in SHANK3-deficent neurons. (A) The PMDS patient-derived ASD04, healthy control PC056, and CRISPR-engineered SHANK3-deficient HT3 and isogenic control WT2 NPC lines were differentiated for 30 days and fluorescently stained. Using quantitative image analysis, in Synapsin 1/SHANK3 double positive synapses located in the MAP2 network the mean SHANK3 signal intensity was quantified. Each datapoint represents technical replicate data from one 384-well plate well. Larger markers represent the medians of independent differentiation experiments. The boxplots show the median and the 2^nd^ and 3^rd^ quartiles of the data. Whiskers indicate 1.5X of the interquartile range. Welch’s unequal variances t-test was performed using the medians of independent differentiation experiments. ns = not significant, * = p < 0.05.

**Figure S3 referring to Figure 4:**
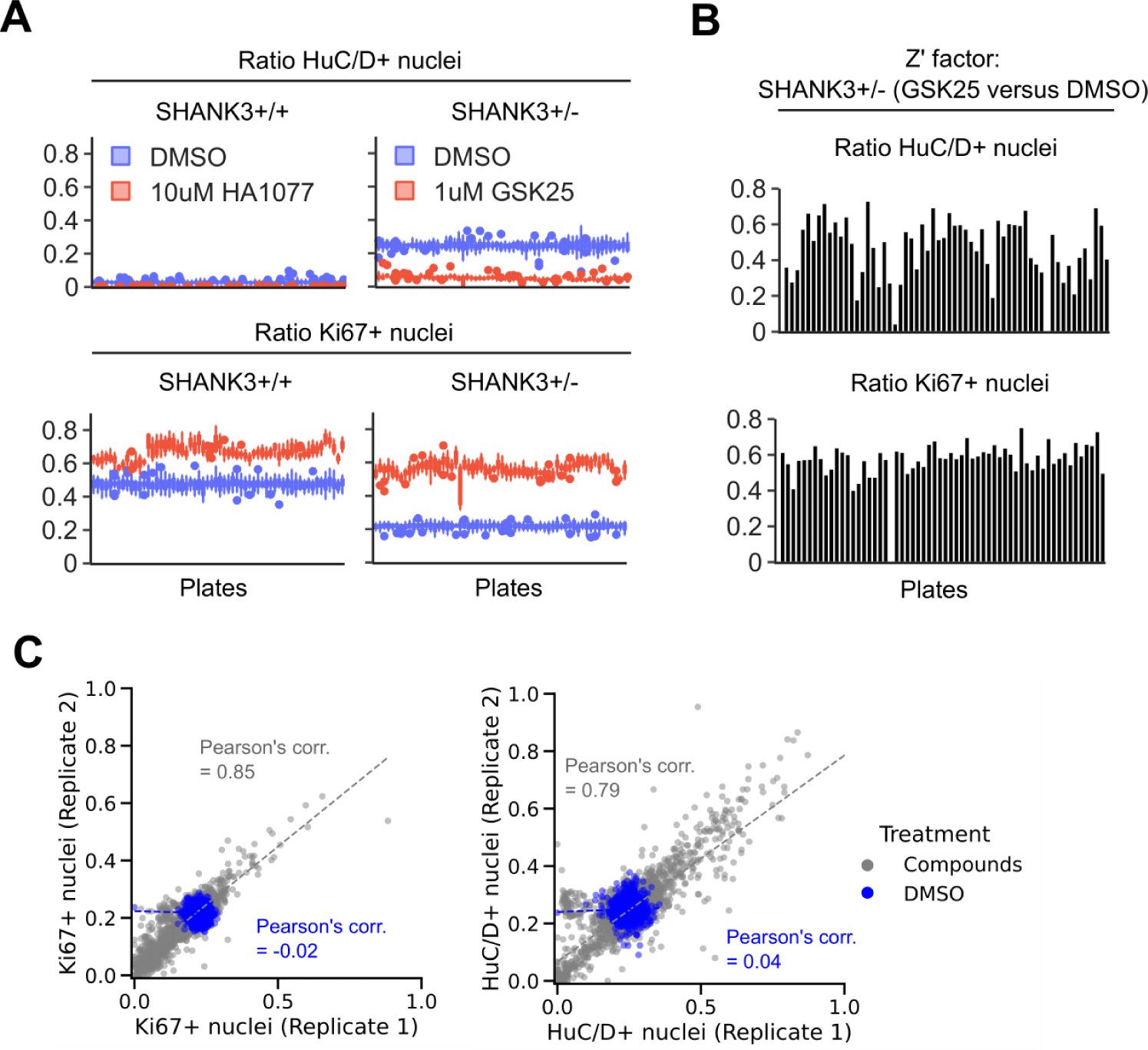
Quality assessment of small molecules screening in CRISPR-engineered NPCs. (A) Ratios of HuC/D and Ki67 positive WT2 and HT3 cells across screening plates. Cells were treated with neutral control DMSO or positive proliferation-inducing control compounds HA1077 (WT2) or GSK-25 (HT3). (B) Z’ factor per plate calculated between chemical controls DMSO and GSK-25. (C) Relationship of HuC/D and Ki67 positivity ratio of compound- and DMSO-treated wells for both replicates. The Pearson correlation coefficients are indicated.

**Figure S4:**
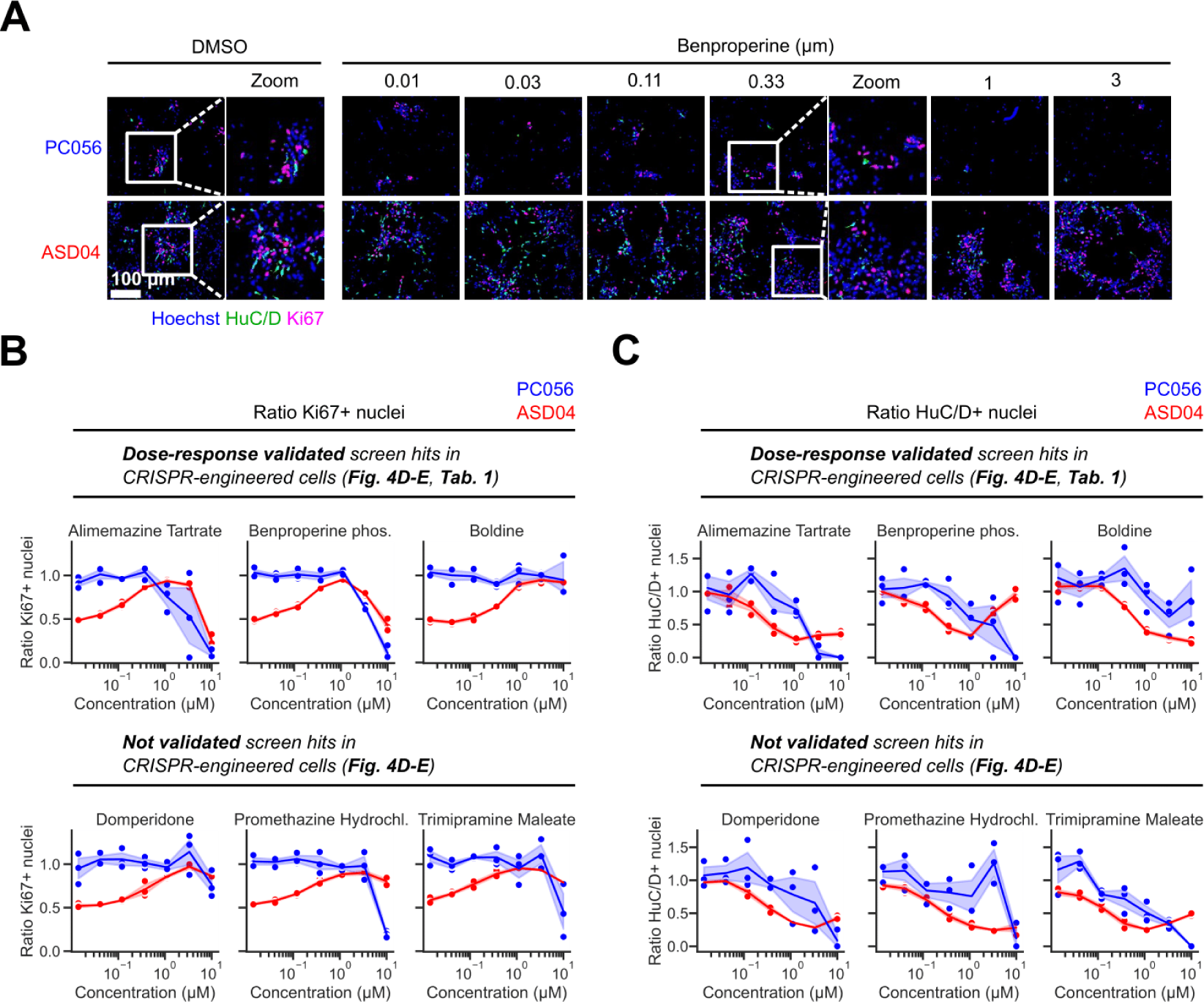
Validation of SHANK3-specific hyperdifferentiation rescuing chemical compounds in PMDS patient and healthy control-derived NPCs. (A) Representative images of healthy control NPC line PC056 and PMDS patient cell line ASD04 stained against proliferation marker Ki67 and differentiation marker HuC/D and treated with DMSO or Benproperine. (B) Dose-response effects of 6 compounds on the ratio of differentiating HuC/D positive ASD04 and control PC056 NPCs. The shaded area in the line graphs corresponds to the 95% confidence interval of triplicate experiments.

**Figure S5 referring to Figure 5:**
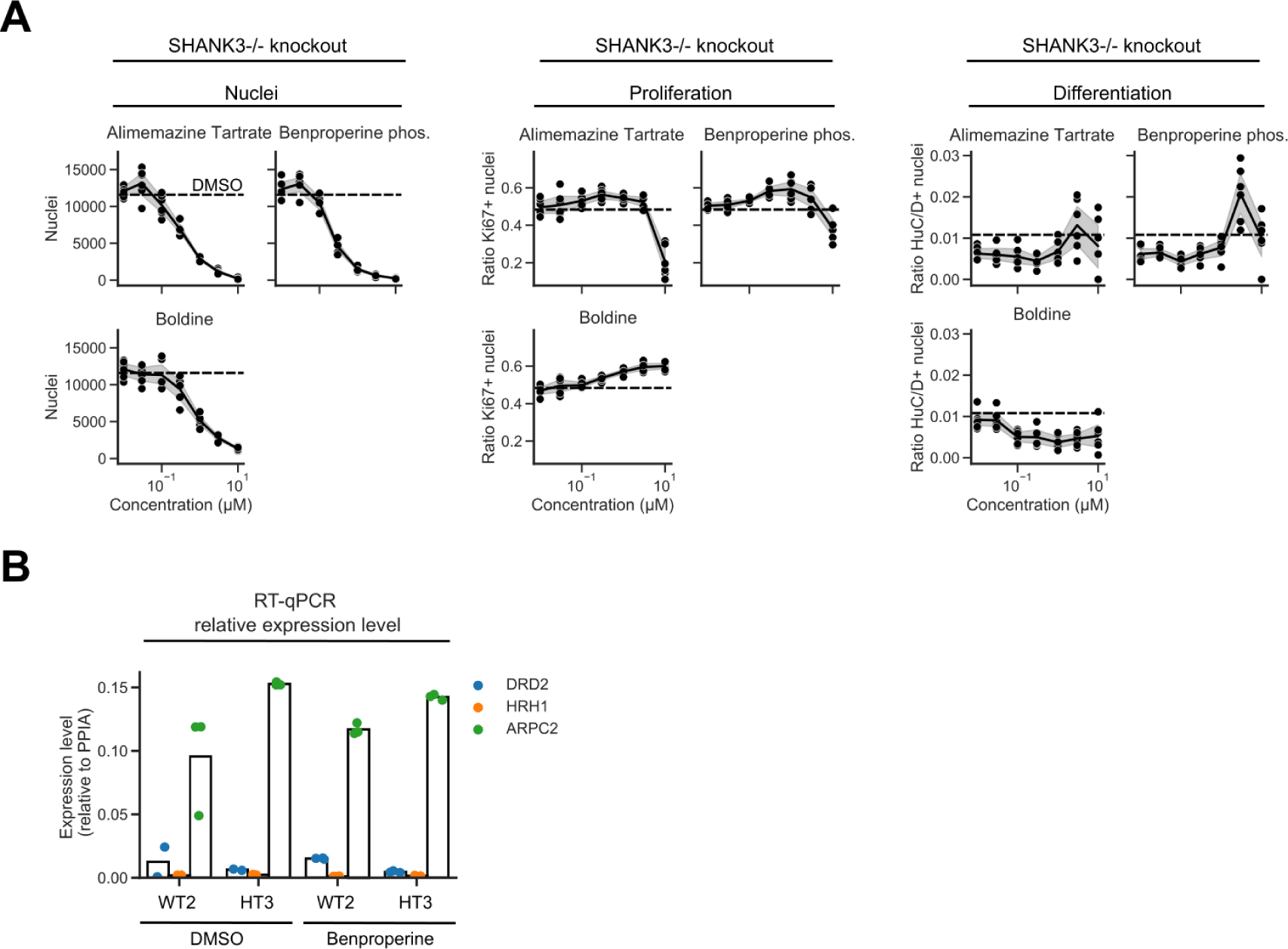
Identified compounds require residual SHANK3 levels for activity. (A) Nuclei count, and ratios of Ki67 and HuC/D positive cells in knockout SHANK3-/-D6 NPCs. The dashed line indicates the DMSO baseline. The shaded area in the line graphs corresponds to the 95% confidence interval. (B) RT-qPCR was performed using Taqman probes against annotated compound targets DRD2, HRH1, and ARPC2 on D6. The expression level was determined relative to PPIA. The individual datapoints of three independent experiments are shown. The bars indicate the median value.

**Figure S6 referring to Figure 6:**
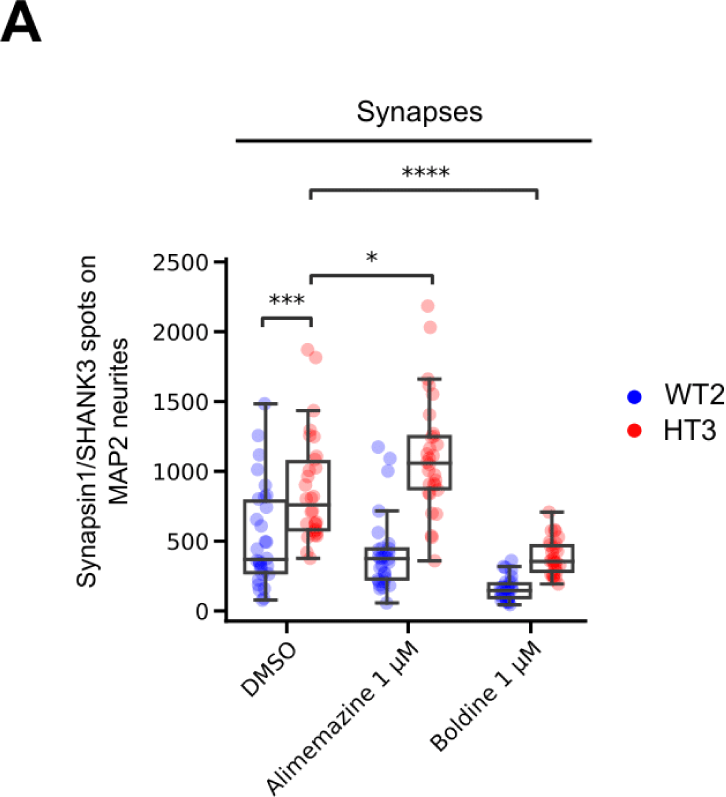
Identified compounds impact synapse formation. (A) HT3 mutant or WT2 isogenic control cells were treated with Alimemazine and Boldine early (D1-3) during differentiation and synapses were quantified after 35 days of differentiation. The boxplots show the median and the 2^nd^ and 3^rd^ quartiles of the data. Whiskers indicate 1.5X of the interquartile range. Welch’s unequal variances t-test was performed. ns = not significant, * = p < 0.05, ** = p < 0.01, *** = p < 0.001.

## Supplementary Tables

**Table S1:** SHANK3-deficiency specific and dose-response confirmed chemical compounds leading to hyperproliferation and hypodifferentiation in CRISPR-engineered NPCs.

| Compound name | CAS number | Target | Detailed annotation (Selleckchem) | SHANK3 specificity in CRISPR NPCs | SHANK3 specificity in patient NPCs |
| --- | --- | --- | --- | --- | --- |
| Alimemazine Tartrate | 4330-99-8 | Unknown | Phenothiazine derivative that is used as an antipruritic. | ++<br>(toxic at high conc. in control) | ++ |
| Benproperine phosphate | 19428-14-9 | Actin-related protein 2/3 complex subunit 2 (ARPC2) | Cough suppressant. Is an orally active, potent actin-related protein 2/3 complex subunit 2 (ARPC2) inhibitor. Attenuates actin polymerization. | ++<br>(toxic at high conc. in control) | ++ |
| Boldine | 476-70-0 | Unknown | Isolated from *Peumus boldus*, has alpha-adrenergic antagonistic properties. Farnesoid X receptor (FXR) agonistic properties. | +++<br>(toxic at high conc. in control) | + |

**Table S2:**
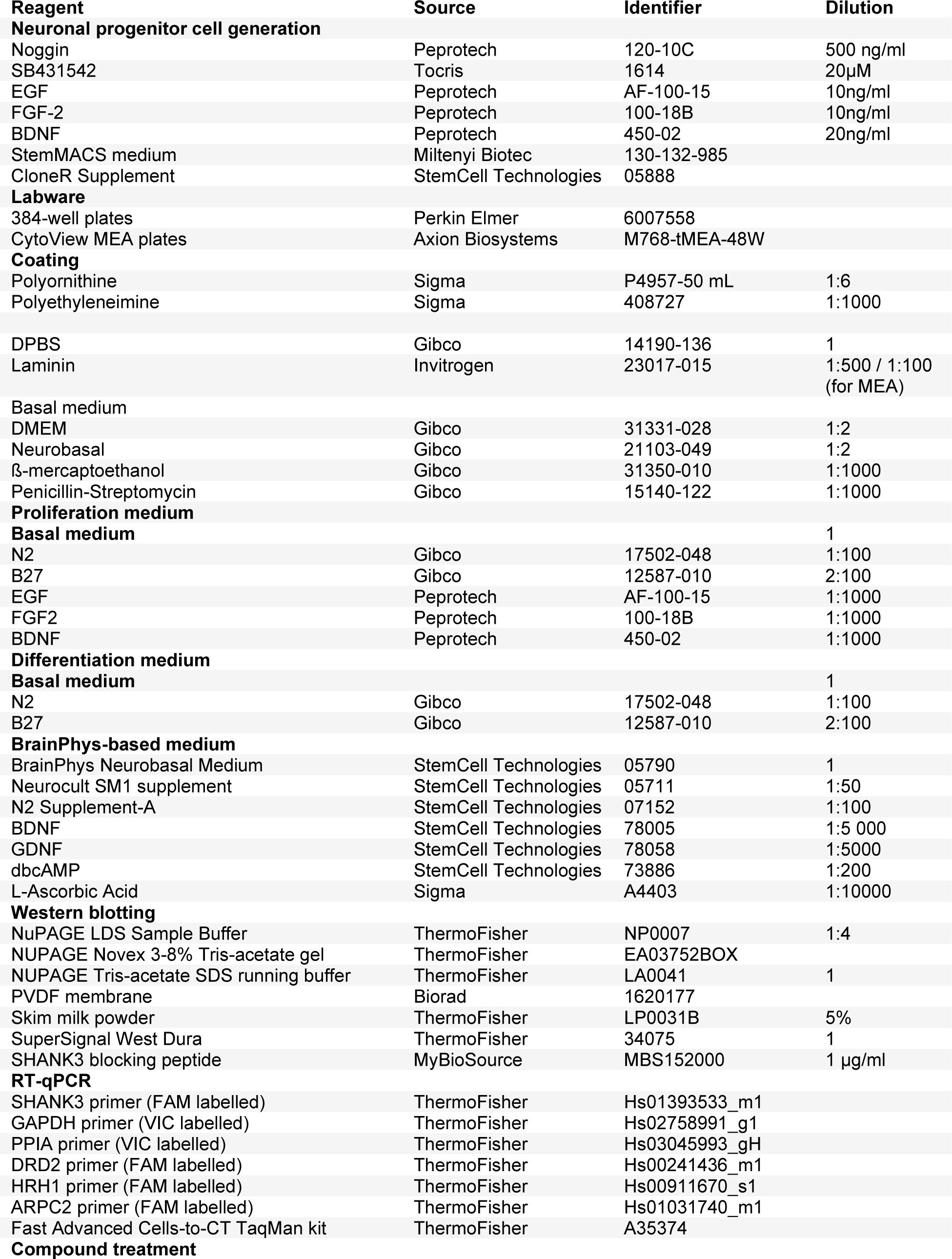
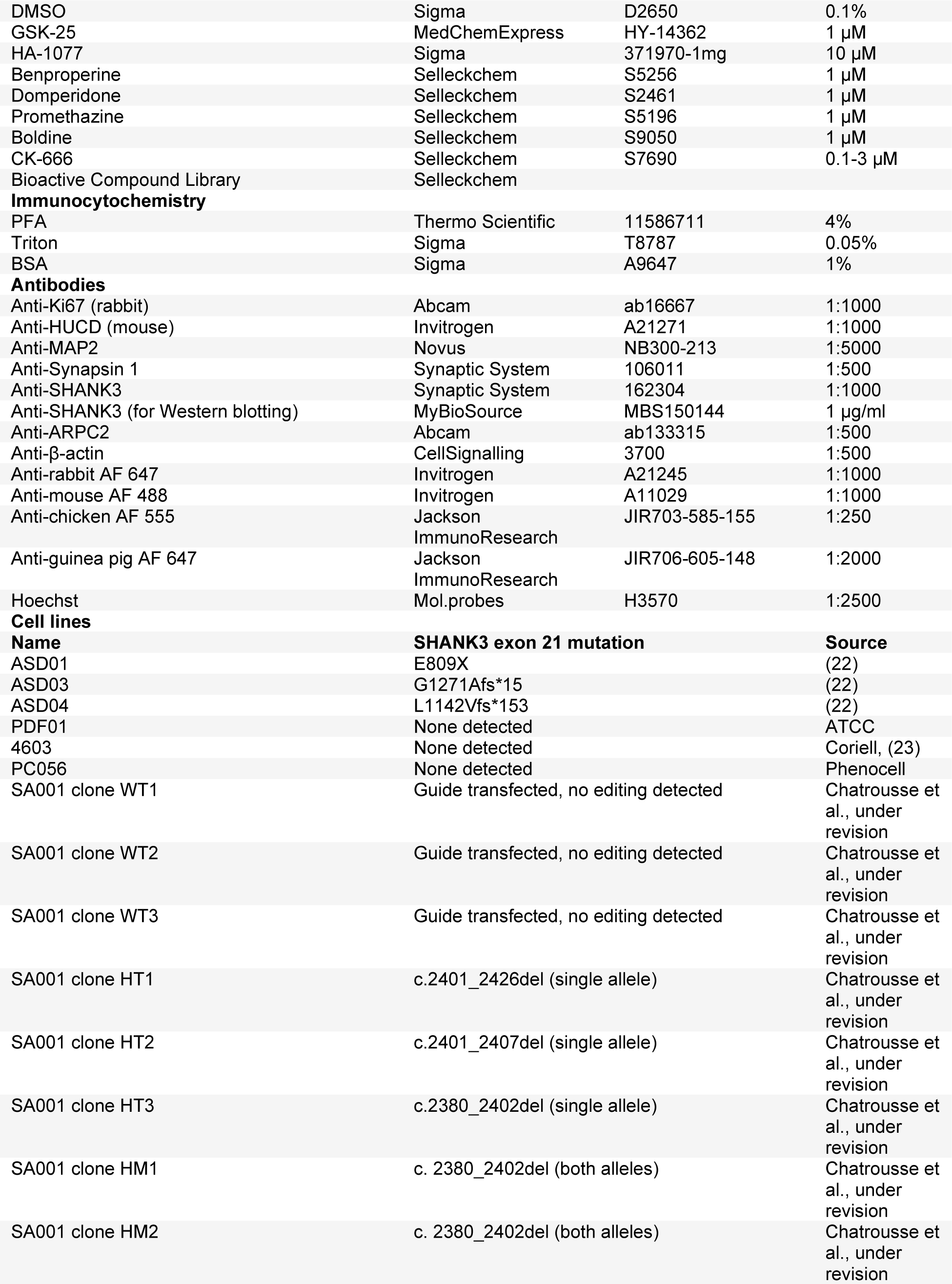
Key resource table.

**Table S3:** Primary screen hits in CRISPR SHANK3+/-neuronal progenitor cell line HT3.

|  | **Compound name** | **CAS number** | **Target** | **Detailed annotation (Selleckchem)** | **Gene names (STRING input)** | **Pathway** |
| --- | --- | --- | --- | --- | --- | --- |
| 1 | A-205804 | 251992-66-2 | Integrin | Inhibitor of E-selectin and ICAM-1 expression with IC50 of 20 nM and 25 nM respectively. | SELE, ICAM1 | Cytoskeletal Signaling |
| 2 | AS2863619 | 2241300-51-4 | CDK | Cyclin-dependent kinase CDK8/19 inhibitor with IC50 of 0.6099 nM and 4.277 nM, respectively. | CDK8, CDK9 | Cell Cycle |
| 3 | AZD1080 | 612487-72-6 | GSK-3 | Selective, orally active, brain permeable GSK3 inhibitor, inhibits human GSK3α and GSK3β with K<sub>i</sub> of 6.9 nM and 31 nM, respectively. | GSK3A, GSK3B | PI3K/Akt/mTOR |
| 4 | AZD2858 | 486424-20-8 | GSK-3 | Selective GSK-3 inhibitor with an IC50 of 68 nM. | GSK3A, GSK3B | PI3K/Akt/mTOR |
| 5 | AZD5363 | 1143532-39-1 | Akt | Inhibits all isoforms of Akt(Akt1/Akt2/Akt3) with IC50 of 3 nM/8 nM/8 nM and lower activity towards ROCK1/2. | AKT1, AKT2, AKT3 | PI3K/Akt/mTOR |
| 6 | AZD5438 | 602306-29-6 | CDK | Inhibitor of CDK1/2/9 with IC50 of 16 nM/6 nM/20 nM in cell-free assays. It also inhibits GSK3β. | CDK1, CDK2, CDK9, GSK3B | Cell Cycle |
| 7 | Alimemazine Tartrate | 4330-99-8 | Others | Phenothiazine derivative that is used as an antipruritic. |  | Others |
| 8 | Autophinib | 1644443-47-9 | Autophagy ,PI3K | Inhibitor with IC50 values of 90 and 40 nM for autophagy in starvation induced autophagy assay and rapamycin induced autophagy assay. The IC50 value for Vps34 is 19 nM in vitro. | PIK3C3 | Autophagy |
| 9 | BI-1347 | 2163056-91-3 | CDK | Inhibitor of CDK8 with IC50 of 1.1 nM. | CDK8 | Cell Cycle |
| 10 | BMS-265246 | 582315-72-8 | CDK | CDK1/2 inhibitor with IC50 of 6 nM/9 nM in a cell-free assay. | CDK1, CDK2 | Cell Cycle |
| 11 | Benproperine phosphate | 19428-14-9 | Others | Cough suppressant. Is an orally active, potent actin-related protein 2/3 complex subunit 2 (ARPC2) inhibitor. Attenuates actin polymerization. | ARPC2 | Others |
| 12 | Boldine | 476-70-0 | FXR receptor | Isolated from Peumus boldus, has alpha-adrenergic antagonist. | NR1H4 | Others |
| 13 | CP21R7 (CP21) | 125314-13-8 | Wnt/beta-catenin | GSK-3β inhibitor that can potentially activate canonical Wnt signalling. | GSK3B | Stem Cells & Wnt |
| 14 | Cepharanthine | 481-49-2 | TNF-alpha | Inhibits tumor necrosis factor TNFα-mediated NFκB stimulation | TNF | Apoptosis |
| 15 | Cerdulatinib<br>(PRT062070,<br>PRT2070) | 1369761<br>-01-2 | JAK | Tyrosine kinase inhibitor with IC50 of 12 nM/6 nM/8 nM/0.5 nM and 32 nM for JAK1/JAK2/JAK3/TYK2 and Syk, respectively. | JAK1, JAK2, JAK3, TYK2, SYK | JAK/STAT |
| 16 | Domperidone | 57808-66-9 | Dopamine<br>Receptor | Oral dopamine D2 receptor antagonist, used to treat nausea and vomiting. | DRD2 | Neuronal<br>Signaling |
| 17 | Doramapimod<br>(BIRB 796) | 285983-48-4 | p38 MAPK | Pan-p38 MAPK inhibitor with IC50 of 38 nM, 65 nM, 200 nM and 520 nM for p38α/β/γ/δ. | MAPK11,<br>MAPK12,<br>MAPK13,MA<br>PK14 | MAPK |
| 18 | GO-203 | 1222186<br>-26-6 | Others | Peptide inhibitor of MUC1-C dimerization. | MUC1 | Others |
| 19 | GSK269962A<br>HCl | 850664-21-0 | ROCK | Selective ROCK inhibitor with IC50 values of 1.6 and 4 nM for ROCK1 and ROCK2, respectively | ROCK1,<br>ROCK2 | Cell Cycle |
| 20 | GSK429286A | 864082-47-3 | ROCK | Selective inhibitor of ROCK1 and ROCK2 with IC50 of 14 nM and 63 nM, respectively. | ROCK1,<br>ROCK2 | Cell Cycle |
| 21 | Geneticin<br>(G418<br>Sulfate) | 108321-42-2 | Anti-<br>infection | Elongation inhibitor of the 80S ribosome. |  | Microbiology |
| 22 | Hexachloroph<br>ene | 70-30-4 | Potassium<br>Channel | KCNQ1/KCNE1 potassium channel activator with EC50 of 4.61 ± 1.29 μM; It can also attenuate Wnt/beta-catenin signaling. | KCNQ1,<br>KCNE1 | Transmembran<br>e Transporters |
| 23 | Isotretinoin | 4759-48-2 | Hydroxylas<br>e | Developed as chemotherapy medication against brain & pancreatic cancer. |  | Metabolism |
| 24 | LY2090314 | 603288-22-8 | GSK-3 | GSK-3 inhibitor for GSK-3α/β with IC50 of 1.5 nM/0.9 nM. | GSK3A,<br>GSK3B | PI3K/Akt/mTOR |
| 25 | Lycorine | 476-28-8 | AChR | Inhibits acetylcholinesterase (AChE) and ascorbic acid biosynthesis. | ACHE | Neuronal<br>Signaling |
| 26 | Lycorine<br>hydrochloride | 2188-68-3 | HCV<br>Protease | HCV inhibitor. |  | Viral Proteases |
| 27 | MSC2530818 | 1883423<br>-59-3 | CDK | CDK8 inhibitor with the IC50 of 2.6 nM. | CDK8 | Cell Cycle |
| 28 | Momelotinib<br>(CYT387) | 1056634<br>-68-4 | JAK | Inhibitor of JAK1/JAK2 with IC50 of 11 nM/18 nM. | JAK1, JAK2 | JAK/STAT |
| 29 | ORY-1001<br>(RG-6016)<br>2HCl | 1431326<br>-61-2 | Histone<br>Demethyla<br>se | Lysine-specific demethylase KDM1A inhibitor with IC50 of <20 nM. | LSD1 | Epigenetics |
| 30 | PHA-767491 | 942425-68-5 | CDK | Cdc7/CDK9 inhibitor with IC50 of 10 nM and 34 nM in cell-free assays, respectively. | CDC7, CDK9 | Cell Cycle |
| 31 | Pexmetinib<br>(ARRY-614) | 945614-12-0 | p38<br>MAPK,Tie-2 | Dual p38 MAPK/Tie-2 inhibitor with IC50 of 4 nM/18 nM. | MAPK11,<br>MAPK12,<br>MAPK13,MA<br>PK14, TEK | MAPK |
| 32 | Promethazine<br>HCl | 58-33-3 | Histamine<br>Receptor | Histamine H<sub>1</sub> receptor antagonist, used as a sedative and antiallergic medication. | HRH1 | Neuronal<br>Signaling |
| 33 | RKI-1447 | 1342278-01-6 | ROCK | Inhibitor of ROCK1 and ROCK2, with IC50 of 14.5 nM and 6.2 nM, respectively. | ROCK1, ROCK2 | Cell Cycle |
| 34 | RepSox | 446859-33-2 | TGF-beta/Smad | Inhibitor of the TGFβR-1 with IC50 of 23 nM. | TGFBR1 | TGF-beta/Smad |
| 35 | SR-4370 | 1816294-67-3 | HDAC | Inhibitor of class I HDACs with IC50 of 0.13 μM, 0.58 μM, 0.006 μM, 2.3 μM, 3.7 μM for HDAC 1, HDAC 2, HDAC 3, HDAC 8, HDAC 6, respectively. | HDAC1, HDAC3 | Epigenetics |
| 36 | Skepinone-L | 1221485-83-1 | p38 MAPK | p38α-MAPK inhibitor with IC50 of 5 nM. | MAPK14 | MAPK |
| 37 | TP0427736 HCl | 864374-00-5 | ALK | Inhibitor of TGFβR-1 with an IC50 of 2.72 nM. | TGFBR1 | Angiogenesis |
| 38 | Tretinoin | 302-79-4 | Retinoid Receptor | Ligand for both the retinoic acid receptor (RAR) and the retinoid X receptor (RXR). | RARA, RARB, RARG, RXRA, RXRB, RXRG | Metabolism |
| 39 | Trimipramine Maleate | 521-78-8 | Others | Antidepressant; serotonin transport blocker that also blocks norepinephrine uptake. |  | Others |
| 40 | UM171 | 1448724-09-1 | Others |  |  | Others |
| 41 | WS6 | 1421227-53-3 | IκB/IKK | Cell proliferation inducer via modulation of Erb3 binding protein-1 (EBP1) and the IκB kinase pathway. | PA2G4, IKBKB | NF-κB |
| 42 | Y-39983 HCl | 173897-44-4 | ROCK | Selective rho-associated protein kinase(ROCK) inhibitor with an IC50 of 3.6 nM. | ROCK1, ROCK2 | Cell Cycle |

**Table S4:** Used cell lines.

| iPSC lines |  |  |  |  |
| --- | --- | --- | --- | --- |
| Cell line name | SHANK3 mutation type | Genetic abnormality | Inheritance | Sex |
| ASD01 | SHANK3 STOP truncation | E809X | de novo | F |
| ASD03 | SHANK3 frame shift truncation | G1271Afs*15 | de novo | M |
| ASD04 | SHANK3 frame shift truncation | L1142Vfs*153 | de novo | M |
| PDF01 | None detected | None detected | N/A | M |
| 4603 | None detected | None detected | N/A | M |
| PC056 | None detected | None detected | N/A | F |
| hESC lines (all clones derived from hESC line SA001) |  |  |  |  |
| Cell line name | Clone | Confirmed CRISPR editing (Shank3 Exon 21) | Predicted effect on translation | Sex |
| WT1 | SA01 g141 cl2 B04 | Guide transfected, no editing detected | N/A | M |
| WT2 | SA01 g141 cl2 H07 | Guide transfected, no editing detected | N/A | M |
| WT3 | SA01 g141 cl3 G04 | Guide transfected, no editing detected | N/A | M |
| HT1 | SA01 g141 cl2 D04 | c.2401_2426del | p.R802Qfs*18 (C-terminal deletion) | M |
| HT2 | SA01 g141 cl3 D05 | c.2401_2407del | p.L801Qfs*54 (C-terminal deletion) | M |
| HT3 | SA01 g141 cl7 A02 | c.2380_2402del | p.A796Rfs*25 (C-terminal deletion) | M |
| HM1 | SA01 g150 cl1 E07 | c. 2380_2402del (both alleles) | p.L801Qfs*49 (C-terminal deletion) | M |
| HM2 | SA01 g150 cl1 C07 | c. 2380_2402del (both alleles) | p.L801Qfs*49 (C-terminal deletion) | M |

